# Charting Tissue Expression Anatomy by Spatial Transcriptome Decomposition

**DOI:** 10.1101/362624

**Authors:** Jonas Maaskola, Ludvig Bergenstråhle, Aleksandra Jurek, José Fernández Navarro, Jens Lagergren, Joakim Lundeberg

## Abstract

We create data-driven maps of transcriptomic anatomy with a probabilistic framework for unsupervised pattern discovery in spatial gene expression data. With convolved negative binomial regression we discover patterns which correspond to cell types, microenvironments, or tissue components, and that consist of gene expression profiles and spatial activity maps. Expression profiles quantify how strongly each gene is expressed in a given pattern, and spatial activity maps reflect where in space each pattern is active. Arbitrary covariates and prior hierarchies are supported to leverage complex experimental designs.

We demonstrate the method with Spatial Transcriptomics data of mouse brain and olfactory bulb. The discovered transcriptomic patterns correspond to neuroanatomically distinct cell layers. Moreover, batch effects are successfully addressed, leading to consistent pattern inference for multi-sample analyses. On this basis, we identify known and uncharacterized genes that are spatially differentially expressed in the hippocampal field between Ammon’s horn and the dentate gyrus.

---

The analysis of spatially stratified, transcriptome-wide data [1–3] poses additional challenges compared to classical analysis of bulk RNA sequencing (RNA-Seq) samples. In the classical setting, annotated covariates dictate how samples are to be grouped and compared. These covariates are typically known, and often controlled for, as is the case when performing differential gene expression (DGE) analysis for bulk sequencing count data in DESeq2 [4]. In contrast, the covariates determining spatial gene expression are often unknown and can change both gradually or abruptly. For example, the number of infiltrating immune cells per unit area often follows smooth gradients, yet tissue boundaries dramatically impact gene expression over small distances.

If the cell types underlying a sample are well characterized, it becomes analytically possible to determine mixing proportions of the cell types’ known expression profiles in spatial gene expression data. But, in the general case, there exists the dual discovery problems of expression profiles and spatial distributions.

The statistical tests Trendsceek and SpatialDE can be used to quantify the extent of spatial variation of individual genes’ expression [5, 6]. Transcending individual genes, the “automatic expression histology” feature of SpatialDE implements a gene clustering approach to hidden pattern discovery. Such clustering formulations, however, appear challenged by genes that participate in multiple, spatially overlapping expression programs, and grade-of-membership formulations seem more appropriate for hidden pattern discovery.

Joint analysis of multiple data sets substantially increases statistical power and sensitivity but is challenging due to the presence of batch effects. Both nonparametric [7, 8] and model-based approaches are used to address such effects in the analysis of single cell RNA sequencing (scRNA-Seq) data. The model-based methods perform regression for the count data based on known sample-level covariates and allow for the discovery of unknown ones.

Log-normal expression models are used for bulk [9, 10] and single cell [11, 12] RNA-seq data. However, they do not faithfully reflect the discrete count nature of RNA-Seq data. Alternatives are discrete count expression models, such as models based on the Poisson [13] or the negative binomial [14, 15] distributions, with the latter being better suited for modeling over-dispersed gene expression data.

ZINB-WaVE [14] offers a zero-inflated negative binomial (ZINB) regression framework for unknown covariate discovery, including gene-level covariates. Embedded in a hierarchical probabilistic ZINB model, scVI [15] utilizes neural networks to model non-linear gene-level responses based on both a latent space and known sample-level covariates.

Interpolating properties of bulk RNA-Seq and scRNA-Seq, Spatial Transcriptomics [3] (ST) count data reflects the gene expression of multiple, but comparatively few cells. Since not all of these cells need to be of the same type, probabilistic models of ST data should admit a mixture interpretation on the level of the counts in each spot.

No method has previously been described to analyze spatial gene expression data that fulfills all of the following criteria: have a discrete count model; be applicable to over-dispersed gene-expression data; able to cope with multiple samples and covariates; admit a mixture interpretation. We outline such a method below.

## Results

Here we describe spatial transcriptome decomposition (STD), a hidden pattern discovery method, which uses convolved hierarchical negative binomial regression to identify transcriptomic patterns in space. We first give an overview of the method before we present several applications with real biological data and evaluates its performance, both on synthetic data and in comparison to related methods.

### Spatial Transcriptome Decomposition

Figure 1 illustrates an application of our method, STD. It performs inference with a probabilistic model of the counts observed for each gene in each spot. Then it computes expected values of marginal relative frequencies: gene expression profiles and spatial activity maps for a set of transcriptomic factors. Like for other model-based methods, these serve as an effective, lower-dimensional representation of the data, and are used for downstream analysis.

**Figure 1:**
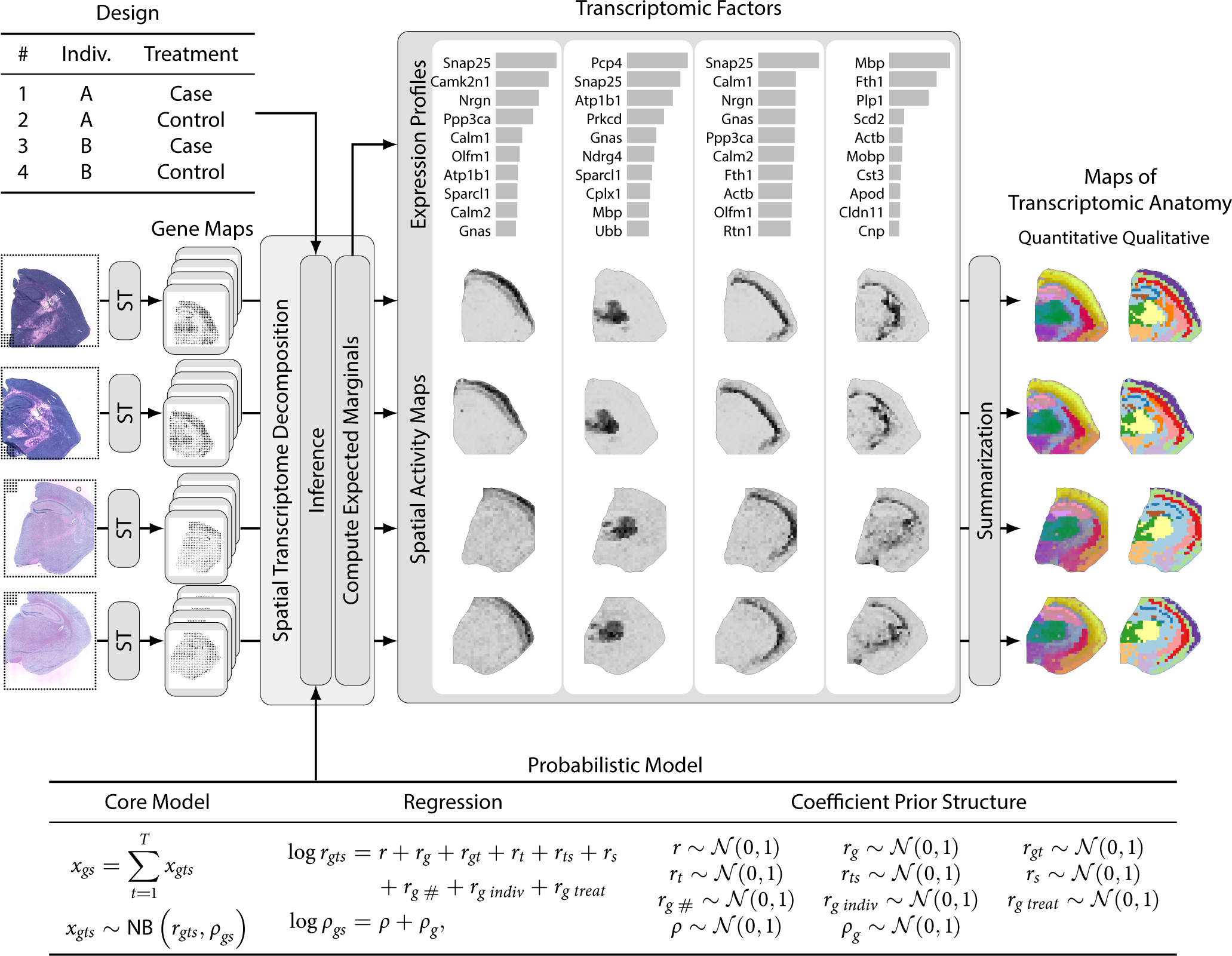
Overview of Spatial Transcriptome Decomposition (STD). Spatial Transcriptomics (ST) is performed for multiple sections from multiple individuals, yielding a set of gene maps, one for each gene and each sample, quantifying the number of reads, *x*_*gs*_, observed for gene *g* in spot *s*. STD performs inference to determine point estimates for all parameters, and subsequently computes expectations of marginal relative frequencies. Thus, the gene maps are decomposed by STD into a set of transcriptomic factors, each comprising one gene expression profile and for each sample a spatial activity map. Expression profiles quantify how strongly each gene is expressed in a given factor, and spatial activity maps reflect where in space each factor is active. Only four factors are indicated, and only ten genes per expression profile. Figure S1 illustrates how Voronoi tesselation is here used to visualize spatial gene expression. In addition to the data, the inference of STD depends on the experimental design and the probabilistic model is adapted to the specific application. The model is composed of three parts: 1) the core model specifying that the observed counts *x*_*gs*_ are the sum of the hidden counts *x*_*gts*_ of several transcriptomic factors *t*, which in turn are negative binomially distributed; 2) the regression equations for the logarithms of the rate and odds parameters of the negative binomial distribution, *r*_*gts*_ and *ρ*_*gs*_; 3) the probabilistic structure specifying the prior distributions of the regression coefficients. The regression equations and prior structure are adapted to the specific application. To illustrate application adaptation, the rate regression equation contains terms *r*_*g* *#*_, *r*_*g* *indiv*_, and *r*_*g treat*_ for gene-dependent terms respectively specific to the sample, individual, and treatment. Finally, the spatial activity maps can be summarized across the factors to yield maps of transcriptomic anatomy, either quantitatively for visual inspection or qualitatively—using hierarchical clustering—for down-stream analyses. Figure S2 illustrates how quantitiative maps of transcriptomic anatomy are created.

In order to leverage increased statistical power, the method is designed for the joint analysis of multiple Spatial Transcriptomics (ST) [3] data sets. On a high level, the inputs to the method are one or more ST count matrices, a description of the experimental design, as well as an adaptation of the probabilistic model to the specific application.

#### Core model

We assume that the observed count *x*_*gs*_ for gene *g* in spot *s* is the sum of hidden counts *x*_*gts*_ due to *T* transcriptomic factors,

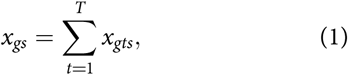

and that these in turn are negative binomially distributed,

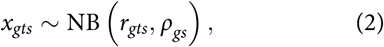

according to eq. (5), with rate and odds parameters *r*_*gts*_ and *ρ*_*gs*_. Notably, the odds parameters *ρ*_*gs*_ are restricted to not depend on the factor (Online Methods).

The choice of the number of factors is a critical parameter of our approach; it determines whether sufficient factors are available or whether factors are over-allocated. This choice is currently left to the user, and it may necessitate some experimenting. In the future, we envisage to employ non-parametric process priors to perform inference across numbers of factors.

#### Regression and experimental design

When jointly analyzing a set of count matrices resulting from multiple ST experiments, both technical and biological variation between the samples needs to be accounted for. To this end, our method performs regression for the negative binomial distribution’s log rate and log odds parameters, with a probabilistic model adapted to the specific application by the analyst. Thus, in addition to the data, the analyst needs to provide the experimental design and to adapt the model to it, by specifying the regression equations as well as the probabilistic prior structure. The framework offers flexible modeling choices for the prior structure: hierarchical probabilistic structures composed from common exponential-family distributions. The graphical probabilistic structure of the model is specified using conventional mathematical notation in model specification files, as described in the Online Methods.

In this communication, we will consider only non-hierarchical, standard normal priors on the regression coefficients.

#### Visualization

Designed in a grid arrangement, the geometries of the ST microarrays exhibit slight irregularities due to technical variability in the printing process. To both accurately represent individual spots’ positions and to address missing data in a visually unobtrusive manner, we utilize Voronoi-tessellation for visualization of inference results (fig. S1).

#### Maps of transcriptomic anatomy

When dealing with spatial maps for numerous features—whether genes or factor activities—it is frequently helpful to condense the information present in them. This can be done with dimensionality reduction techniques from machine learning, such as t-distributed stochastic neighbor embedding (t-SNE) [16], principal component analysis (PCA), or uniform manifold approximation and projection (UMAP) [17]. By applying such a technique, we compress the information across all features into three components. After rescaling into the unit cube, we use these as coordinates in color space to colorize the spots in spatial plots. Feature similarity is thus encoded by colors, and when the features reflect transcriptome-wide gene expression data we refer to such plots as maps of transcriptomic anatomy (fig. S2).

### Mouse olfactory bulb

We jointly analyze sixteen mouse olfactory bulb ST libraries from seven individuals, including four previously-unpublished libraries (table T1 and Online Methods).

**Table T1:** Design of mouse olfactory bulb samples. Note that the first twelve samples were previously published by Ståhl et al. and that individual B and female sex are confounded.

| Section | Individual | Sex | Reference |
| --- | --- | --- | --- |
| 1 | A | ♂ | Ståhl et al. [1] |
| 2 | A | ♂ | Ståhl et al. [1] |
| 3 | A | ♂ | Ståhl et al. [1] |
| 4 | A | ♂ | Ståhl et al. [1] |
| 5 | B | ♀ | Ståhl et al. [1] |
| 6 | C | ♂ | Ståhl et al. [1] |
| 7 | C | ♂ | Ståhl et al. [1] |
| 8 | D | ♂ | Ståhl et al. [1] |
| 9 | D | ♂ | Ståhl et al. [1] |
| 10 | E | ♂ | Ståhl et al. [1] |
| 11 | E | ♂ | Ståhl et al. [1] |
| 12 | E | ♂ | Ståhl et al. [1] |
| 13 | F | ♂ | here |
| 14 | F | ♂ | here |
| 15 | G | ♂ | here |
| 16 | G | ♂ | here |

The factor analysis was performed for 20 factors utilizing staging, 20% dropout frequency, and adaptive down-sampling to equate sequencing depth (Online Methods). Results for a subset of samples and factors are shown in fig. 2; fig. S5 displays results for all samples and factors.

**Figure 2:**
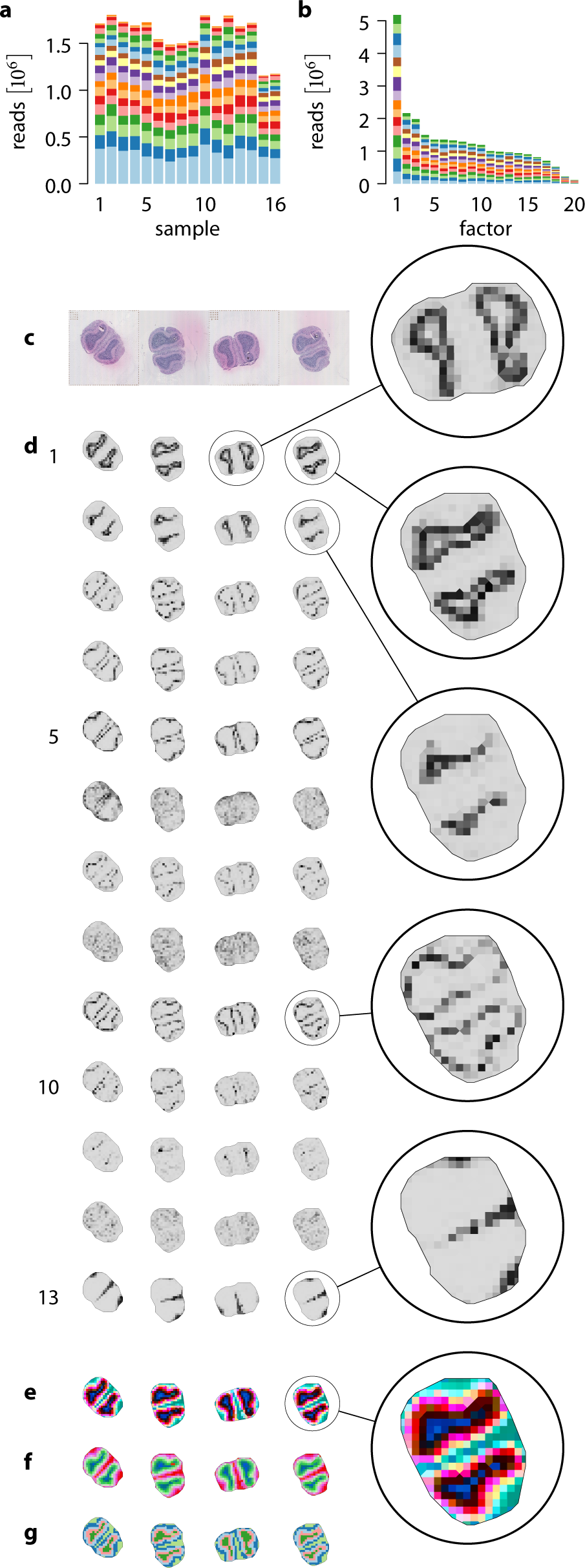
Transcriptomic patterns in mouse olfactory bulb sections. (**a**) Number of reads across samples, colors indicate factor. (**b**) Number of reads across factors, colors indicate sample. (**c**) H&E-stained microscopy images for 4 of 16 sections. All sections are shown in fig. S5. (**d**) Spatial factor activity maps for 13 of 20 factors. All factors are shown in fig. S5. (**e**, **f**) Summarization of spatial factor activities, using t-SNE (**e**) or UMAP (**f**). (**g**) Hierarchical clustering of factor activities into five clusters.

Sequencing depth is comparable across the samples (fig. 2a), and the relative proportions of reads and spots attributed to the different factors are approximately constant across the samples (figs. S3c and S3d). Factors are ordered by decreasing number of attributed reads, and top-ranking factors have markedly more reads attributed than lower ranking ones (fig. 2b).

Microscopy images of the H&E-stained cryo-sections and inferred spatial factor activity maps are shown in figs. 2c and 2d. The spatial activities of most inferred factors reflect the olfactory bulb anatomy across individuals and replicates, while for four factors the correspondence to anatomy is less clear (factors 6, 8, 12, 20 in figs. 2d and S5b). Summarizing the spatial factor activities by t-SNE or UMAP (figs. 2e and 2f) yields colorizations of the spots consistent with anatomical position across individuals and replicates. But when the same summarization techniques are applied directly to the read counts (figs. S6 to S8), rather than to the spatial factor activities, then uncorrected effects are visible between samples that impede identification of corresponding regions across individuals and replicates.

We partitioned the spots into five sets based on hierarchical clustering of the spatial factor activities (figs. 2g and S5e). From outside inwards, these clusters correspond to the olfactory nerve layer, the glomerular layer, the plexiform and mitral cell layers, as well as two for the granular cell layer: a peripheral and a central one. The outer plexiform, mitral, and inner plexiform layers cannot clearly be resolved by partitioning into more clusters (results not shown), presumably due to insufficient spatial resolution of the array spots. Applying hierarchical clustering directly to the read counts (fig. S9), rather than to the spatial factor activities, again exhibits uncorrected between-sample effects.

Subsequently, we performed DGE analysis using DESeq2 between all pairs of clusters (Online Methods). We intersected the sets of genes that are significantly up-regulated in a given cluster in all pairwise analyses (supplementary dataset D1) to define sets of genes that are specific to that cluster. We then retrieved images of in situ hybridizations (ISH) for the cluster-specific genes from the Allen brain atlas [18]. Inspection of representative ISH images (fig. S10) reveals that the cluster-specific genes share common, cluster-specific spatial expression patterns in the Allen brain atlas.

### Coronal brain sections

We analyze four ST libraries prepared from mouse coronal brain sections, neighboring sections from two individuals. These sections contain parts of the hippocampus, cortex, cerebral nuclei, thalamus, hypothalamus, as well as several cross-cutting fiber tracts.

Like for the olfactory bulb data, the factor analysis was again performed for 20 factors utilizing staging, 20% dropout frequency, and adaptive down-sampling to equate sequencing depth (Online Methods). Results for all samples and factors are shown in fig. 3, and read and spot statistics across samples and transcriptomic factors are displayed in fig. S4.

**Figure 3:**
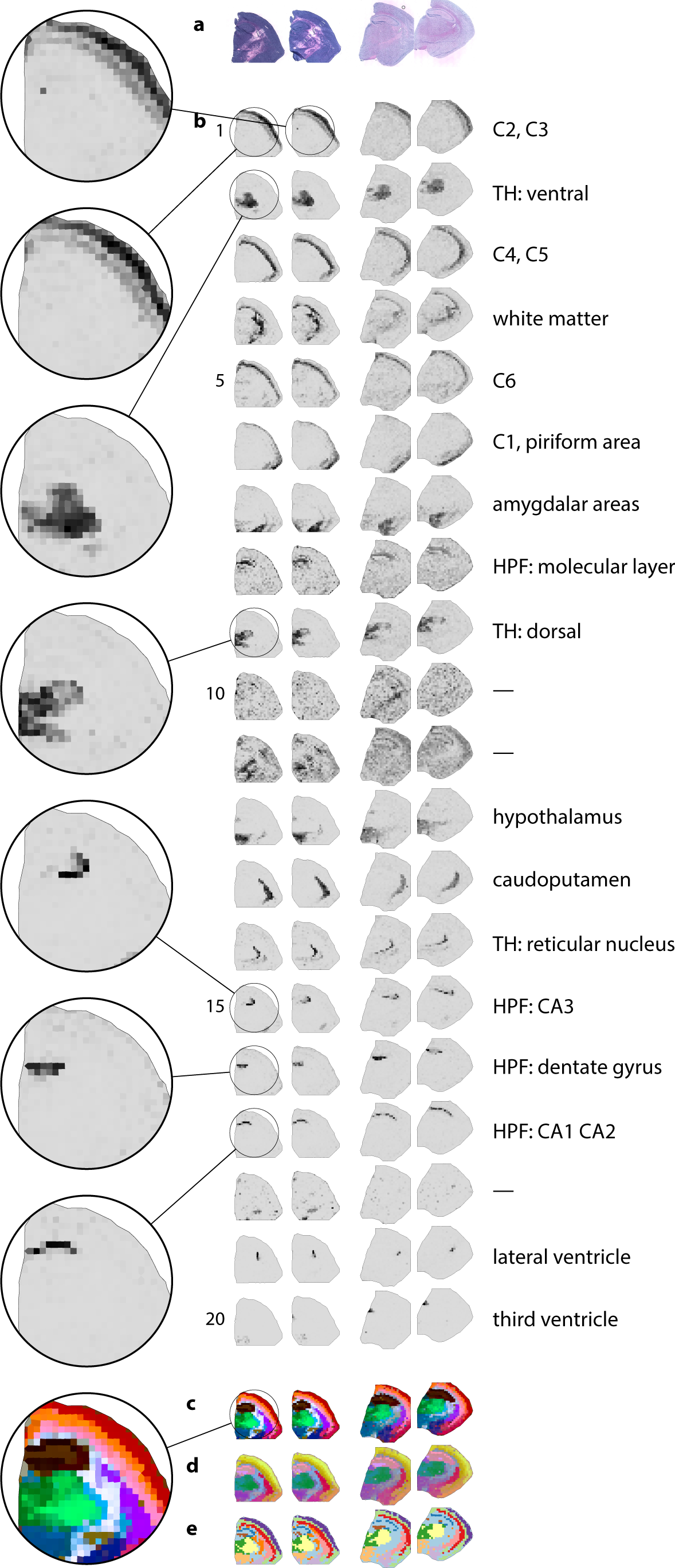
Transcriptomic patterns in mouse coronal brain sections. (**a**) H&E-stained microscopy images. (**b**) Spatial factor activity maps, and neuroanatomical regions co-incident with factor activity. (**c**, **d**) Summarization of spatial factor activities, using t-SNE (**c**) or UMAP (**d**). (**e**) Hierarchical clustering of factor activities into twelve clusters. Abbreviations: HPF hippocampal field, TH thalamus, C1–C6 cortical layers 1–6, CA1–CA3 Ammon’s horn regions 1–3.

Inspection of the spatial factor activity maps reveals that at least 17 of the 20 factors correspond to neuroanatomical brain structures (fig. 3b). In particular, several factors correspond to cortical areas; one each for layer 1, for layers 2 and 3, for layers 4 and 5, and for layer 6, for the striatum-like amygdalar nuclei and for the cortical olfactory areas. Four structures are identified within the hippocampal field: the pyramidal cell layer in Ammon’s horn (one factor for CA1 and CA2, another for CA3), the granule cell layer in the dentate gyrus, and the molecular layer. Thalamic structures are represented by three factors, for the dorsal thalamus, the ventral thalamus, and for the reticular nucleus. Further factors correspond to the hypothalamus, the caudoputamen, the lateral ventricle, the third ventricle, and white matter.

The remaining three factors appear to capture residual signal, as they exhibit more diffuse spatial activities and do not capture the same anatomical structures in both individuals. Factors 11 and 18 capture specific but different anatomical structures within the individuals, and factor 10 competes with the white matter factor in one sample.

Summarizing the spatial activity maps by t-SNE or UMAP consistently colorizes spots in corresponding anatomical positions within and across individuals and replicates (figs. 3c and 3d). Similarly, hierarchical clustering of the spatial activities into twelve clusters consistently partitions the spots into distinct, neuroanatomically-defined regions (figs. 3e and S9d).

We performed pairwise DGE analyses using DESeq2 for all pairs of clusters (supplementary dataset D2). Functional enrichment analysis (Online Methods) for these DGE results yields almost exclusively meaningful categories from the various ontologies (supplementary dataset D3), such as “calcium ion-binding” (GO-MF), “dendrite” and “axon” (GO-CC), “chemical synapse transmission” (GO-BP), “dopaminergic synapse” and “glutamatergic synapse” (KEGG 2016), “neuronal system” (Reactome 2016), among others.

Considering one such pairwise comparison closer, we find 350 genes differentially expressed (adjusted *p*-value < 0.01) between the dentate gyrus and Ammon’s horn clusters, of which 233 are expressed higher in dentate gyrus (fig. 4a). “Dentate gyrus” and “Field CA1, stratum oriens” are the categories from the “Allen_brain_up” ontology that are respectively most enriched for genes significantly differentially regulated in this comparison (adjusted *p*-values 5.7×10^−21^and 3.8×10^−18^). In situ hybridizations from the Allen brain atlas constitute further orthogonal validation of the spatial specificity for the most differentially regulated genes from this comparison (figs. 4b to 4e). Three of these are well-studied proteins with known neuro-biological functions: the transcription factors Prox1 and Neurod6, and the alternative splicing factor Khdrbs3. Our results and the Allen brain atlas suggest that also the predicted membrane protein Fam163b, which has not previously been studied further, might be differentially involved in processes between dentate gyrus and Ammon’s horn.

**Figure 4:**
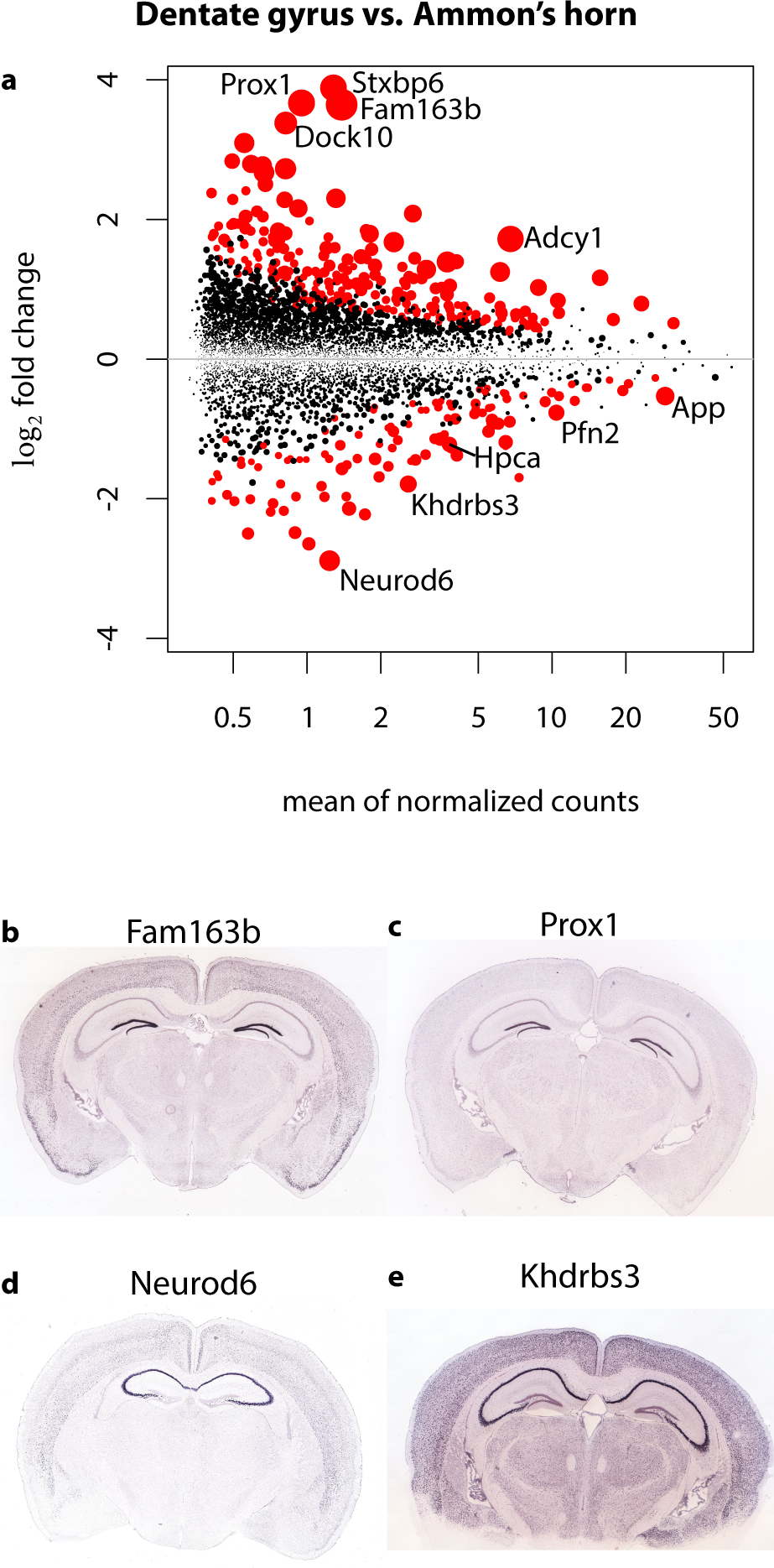
Differential gene expression analysis between the clusters corresponding to dentate gyrus and Ammon’s horn. (**a**) Expression log fold change of dentate gyrus over Ammons horn as a function of mean expression. Top 5 most significantly up- and down-regulated genes are labeled. Circle area is proportional to negative log *p*-value, *p*-value < 0.01 marked red. (**b**–**e**) In situ hybridization images from the Allen mouse brain atlas of genes significantly differentially expressed between Ammon’s horn and dentate gyrus.

### Transcriptomic maps mirror anatomy

Juxtaposition of summarized spatial factor activities with the corresponding region in the Allen mouse brain reference atlas (figs. 5a, 5b, 5d, and 5e) reveals that the resulting spot colorization highlights the same gross anatomical features. This is noteworthy because the reference atlas is based on manually-curated expert knowledge while our results are based on automatic analysis of transcriptome-wide data and do not incorporate any prior knowledge.

**Figure 5:**
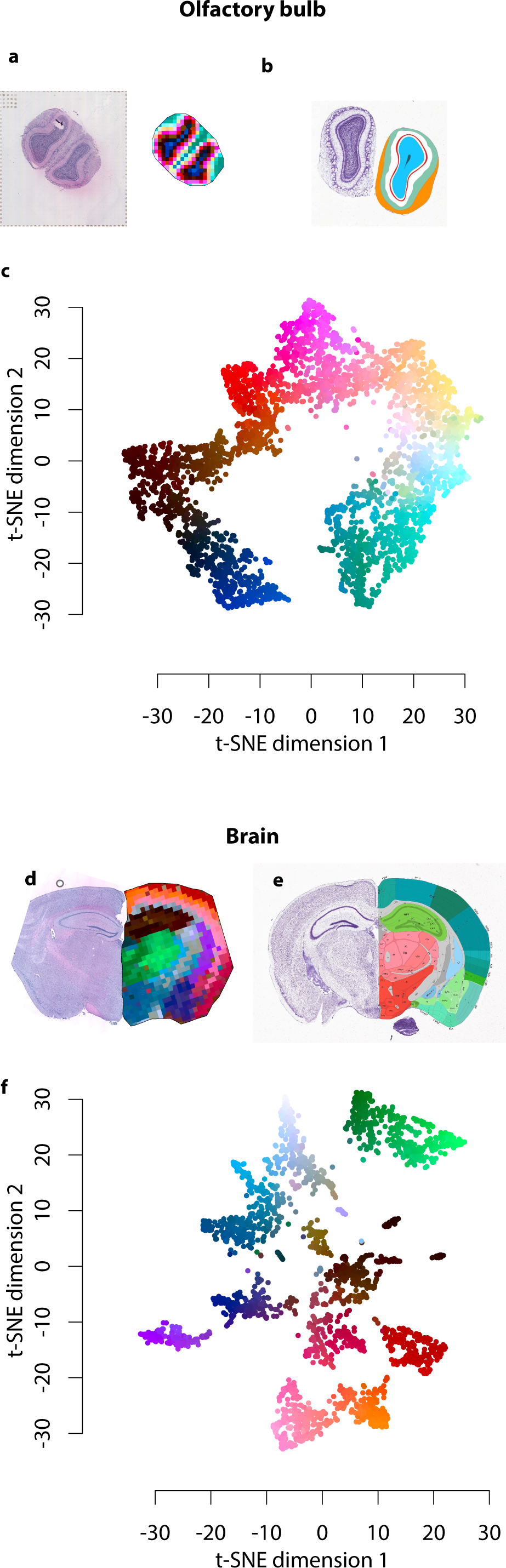
Summarization of factor activities and curated reference atlas. Transcriptomic patterns (**a**, **d**) and corresponding regions in the Allen mouse brain reference atlas (**b, e**) in mouse olfactory bulb (**a**–**c**) and brain sections (**d**–**f**). (**c, f**) Two-dimensional t-SNE summarization of factor activities; colors are identical to (**a, d**) and figs. 2e and 3c, respectively. The mouse olfactory bulb model exhibits a one-dimensional manifold topology corresponding to the inside-outside axis.

Further reducing the dimensionality of the spatial factor activities to two dimensions reveals that the olfactory bulb data is modeled by a topologically one-dimensional manifold (fig. 5c), while the brain data exhibit discrete islands corresponding to disjoint clusters of spots (fig. 5f).

### Performance evaluation

#### Synthetic data

To study the sensitivity of the model to properties of the input data, we simulate synthetic data based on different sets of ground truth parameters. For each such set, the simulated data is decomposed and the result of the decomposition is compared to the ground truth (fig. 6a). We evaluate the success of the decomposition by computing the Pearson correlation between the expected number of reads generated by the true and inferred models in the gene-factor and spot-factor marginals. Details of the data generation and performance evaluation are described in the Online Methods.

**Figure 6:**
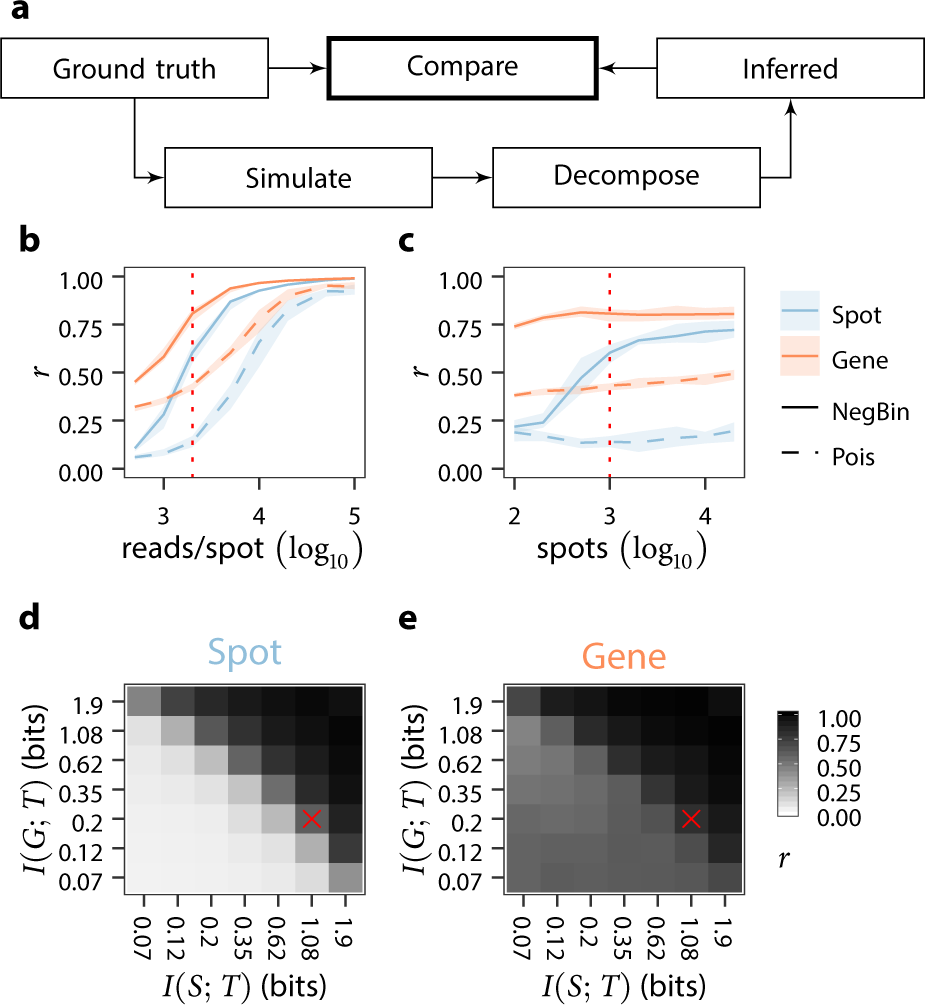
Supervised performance evaluation on synthetic data. (**a**) Based on a set of ground truth parameters, we simulate synthetic expression data. The simulated data is decomposed and the inferred parameters are compared to the ground truth. (**b**–**e**) We run 10 simulations for each unique set of ground truth parameters and investigate model performance along four dimensions: the number of reads per spot, the number of spots, and the heterogeneity of activity maps and gene profiles. The two latter quantities are measured as the mutual information, *I*, between the random variables *S* and *T*, and *G* and *T*, defined so that *p*(*G*=*g, T*=*t, S*=*s*) is the relative frequency of reads for a given gene *g*, transcriptomic factor *t*, and spot *s*. Baselines are representative of real data and annotated as red crosses or dotted lines. (**b, c**) Correlation of expected read number between true and inferred models over the spot-factor and gene-factor marginals; as a function of the number of reads per spot (**b**), or of the number of spots (**c**). Lines correspond to medians and shaded areas to minima and maxima across 10 simulations. Solid lines show results for negative binomial decomposition, as described in this paper, while dashed lines show results for the Poisson regression framework of Berglund et al. [13]. (**d, e**) Median correlation of expected read number between true and inferred models in the spot-factor (**d**) and gene-factor (**e**) marginals across different levels of factor heterogeneity in the same dimensions.

Overall, the performance of the model is positively associated with and highly sensitive to the number of reads (fig. 6b) and the heterogeneity of the transcriptomic factors (figs. 6d and 6e), both in terms of their spatial activities and gene profiles. In contrast, the performance is less sensitive to the number of spots in the input data (fig. 6c). Crucially, for parameter values inferred from real data (annotated in figs. 6b to 6e), the data is decomposed with high accuracy.

#### Comparison with related methods

We first assess the performance of the Poisson regression framework of Berglund *et al.* [13] on the synthetic data described above. Our method shows substantially better performance across all properties of the input data (figs. 6b and 6c). Moreover, this result holds when using a Poisson source model similar to their inference model (Online Methods, fig. S11).

We further assess the performance of two other related methods, scVI [15] and ZINB-WaVE [14], on the brain and olfactory bulb datasets analyzed in this manuscript (Online Methods). Based on a visual comparison of summarized analysis results (figs. S12 and S13), we find both scVI and ZINB-WaVE useful for the analysis of ST data, but less successful than STD at addressing batch effects. The full analysis results of scVI and ZINB-WaVE are visualized in figs. S14 to S17 and discussed in the Online Methods, where we also explain how the empirical observations relate to theoretical differences between the methods.

## Discussion

We presented a probabilistic method to model spatial gene expression count data by convolved negative binomial regression and thereby simultaneously infer unknown gene expression profiles and their unobserved mixing proportions.

The method is related to and generalizes the negative binomial regression models of DESeq2 and ZINB-WaVE, although it does not currently implement a zero-inflation model. While it is quite possible to augment our model accordingly, it is conceivable that our mixture approach may constitute a modeling alternative for zero-inflation. Further, instead of relying on DESeq2 for DGE analysis, it appears worthwhile to investigate the benefits of directly performing DGE analysis based on the parameters inferred by our model. Another relevant future addition to our method might be to allow for Gaussian processes to be used as prior distributions for the spatial coefficients specified in the regression equations. This could help to discover spatially smoother distributions. Another avenue for future improvement is to improve scalability to large data sets. To date, we have successfully applied our method to datasets approaching 10^5^ spots, but data set sizes are expected to grow substantially in the future.

The applications to the olfactory bulb and brain sections demonstrated that our method identifies anatomical regions from the spatial, transcriptome-wide data alone, without requiring additional prior knowledge. Furthermore, the identified patterns are consistent across neighboring sections and individuals, and this consistency indicates that our method successfully corrects technical batch effects. Importantly, when dimensionality reduction or hierarchical clustering are applied directly to gene expression data—rather than to our spatial factor activities—these un-corrected batch effects are evident and may confound down-stream analyses. Automatically and consistently identifying corresponding regions in multiple samples, whether across replicates or across biological contrasts, is a crucial requirement to benefit from sample size increases in downstream statistical analyses.

The Allen mouse brain atlas project provided orthogonal validation for the spatial patterns revealed by our analyses, both in terms of functional enrichment results and in terms of in situ hybridization imaging data for individual genes found differential in our DGE analyses.

To conclude, the spatial factor activity maps inferred by our method quantitatively reflect the spatial expression profiles of corresponding cell types and tissue anatomy. As such they constitute an explanatory interpretation for patterns observed across thousands of genes and provide a foundation for down-stream analyses. Finally, condensing the activity maps of all factors offers a data-driven way to create qualitative and quantitative maps of transcriptomic anatomy.

## Supporting information

Supplementary Dataset D1 - Pairwise DGE analyses - MOB

Supplementary Dataset D2 - Pairwise DGE analyses - Brain

Supplementary Dataset D3 - Pairwise DGE analyses - Brain - Functional Enrichment

## Software availability

The method is implemented in C++ and available as free software from https://github.com/SpatialTranscriptomicsResearch/ std-nb under the terms of the GNU GPL v3 license. Data and results are available in the same place. Additionally, we provide implementations of STD based on PyTorch and TensorFlow that are faster than the C++ version but provide less modeling flexibility.

## Acknowledgements

We thank Tyler Demarest, Vilhelm Bohr, and Deborah Croteau for providing samples. This work was supported by the Knut and Alice Wallenberg Foundation, Swedish Foundation for Strategic Research, the Swedish Research Council, and Science for Life Laboratory. We thank the National Genomics Infrastructure (NGI), Sweden for providing infrastructural support.

## Ethical permits

All work associated with this project has been done in accordance with the National Institute on Aging Animal Care and Use Committee and according to animal study protocol number 361-LMG-2020.

## Contributions

J.M. designed and implemented the method, designed the experiments, and wrote the manuscript. L.B. contributed to the implementation and designed and performed experiments. A.J. performed the wet-lab work. J.F.N. mapped sequencing reads to the genome and quantified genes in each spot. J.La. and J.Lu. supervised the project.

## Online Methods

### Materials

#### Slides with spatially barcoded arrays

To generate the Spatial Transcriptomics data, Codelink Activated Slides (Surmodics) with 1007 distinct capturing oligonucleotides attached were used [1]. Briefly, the oligonucleotides comprising T7 RNA polymerase promotor sequence, 18-mer unique barcode, 9-mer semi-randomized or 7-mer randomized UMI and a poly20TVN capture region were immobilized in 100 µm spots with center-to-center distance of 200 µm. Six 6200 µm × 6600 µm subarrays were printed onto one glass slide.

#### Tissue collection and sectioning

Adult C57BL/6 mice were sacrificed and the brains were removed from the cranial cavity. Olfactory bulbs were dissected out on ice, snap-frozen in dry ice/isopentane slurry and then embedded in OCT (Sakura). The left hemisphere was put into a mold filled with cold OCT and snap-frozen in isopentane pre-cooled with liquid nitrogen. Olfactory bulbs and the left hemisphere were sectioned on the cryostat at 10 µm thickness. Sections were placed on the spatially barcoded arrays with 1 section per well.

#### Fixation, staining and imaging

Sections were fixed for 10 min in 3.6 % to 3.8 % formaldehyde (Sigma) in PBS, washed, then treated for 1 min with isopropanol and air-dried. To stain the tissue, sections were incubated in Mayer’s Hematoxylin (Dako) for 7 min and then Bluing buffer (Dako) for 2 min, followed by Eosin (Sigma) for 20 s. After drying, slides were mounted with 85 % glycerol and imaged using Metafer Slide Scanning Platform (Metasystems). Raw images were stitched together using VSlide software (Metasystems).

#### Tissue permeabilisation

To separate six subarrays from each other and to create reaction chambers for individual sections, the slide was placed in an ArrayIT hybridization cassette. To pre-permeabilize the tissue, sections of olfactory bulbs were incubated for 30 min at 37 °C with Exonuclease I Reaction Buffer (NEB) mixed with 0.2 µg/ µL BSA (NEB). Sections from the hippocampal region were incubated for 20 min at 37 °C with 0.2 U / µL collagenase (Thermo Fisher Scientific) in HBSS buffer (Thermo Fisher Scientific) supplemented with 0.2 µg/ µL BSA. After washing in 0.1x SSC buffer (Sigma), sections of olfactory bulbs and the hippocampal region were permeabilized with 0.1 % pepsin/HCl (Sigma) at 37 °C for 10 min and 6 min, respectively. Then, wells were carefully washed with 0.1x SSC buffer.

#### Reverse transcription and library preparation

Following permeabilisation, reverse transcription mix was added to each well and incubated overnight at 42 °C as described previously [1]. Next, tissue was removed and the surface probes with bound mRNA/cDNA were cleaved from the slide [1]. 65 µL of the reaction mixture containing the released probes were collected from each well and 2nd strand synthesis, cDNA purification, in vitro transcription, aRNA purification, adapter ligation, post-ligation purification, a second 2nd strand synthesis and purification were carried out using an automated MBS 8000 system as described previously [2]. Purified cDNA was then PCR amplified using Illumina Indexing primer [1]. The indexed libraries were purified using carboxylic acid beads on an automated MBS robot system [3] and eluted in 20 µL Elution buffer (Qiagen). The length distribution of the libraries was determined by using the DNA HS Kit (Agilent) with Bioanalyzer 2100 according to the manufacturer’s protocol. The concentration of the libraries was measured with Qubit dsDNA HS (Thermo Fisher Scientific) according to the manufacturer’s protocol. The finished libraries were diluted to 4 nm with Elution buffer and sequenced on the Illumina Nextseq platform using paired-end sequencing, according to the manufacturer’s protocol.

#### Staining of the slide spots and image alignment

After cleaving of the probes from the glass surface, slide was incubated with hybridisation mixture containing Cyanine-3 labelled probes, as described previously [1]. Fluorescent images were acquired using the same scanning platform as for the bright field images. Bright field images and corresponding fluorescent images were aligned manually using Adobe Photoshop CS6 and the spots located under the tissue were selected.

## Bioinformatics

Sequenced reads were processed with the ST Pipeline [4, version 1.4.5] in order to obtain matrices of counts where each cell represents the number of unique molecules for a given spot and a given gene. Homopolymer stretches of at least 10 bp and low quality bases (phred-33 score ≤ 20) were removed from R2. Reads were discarded if R2 was shorter than 20 bp. A contaminant filter was applied to the remaining reads using the Ensembl GRCm38 (v.86) non-coding RNA reference. Filtered reads were then mapped to the genome Ensembl GRCm38 (v.86), demultiplexed and annotated using the reference Mouse GenCode vM11 (Comprehensive gene annotation). Unique counts (UMIs) for each spot/gene combination were computed with default settings of the ST Pipeline. The obtained matrices of counts were processed to replace Ensembl ids by gene names where only protein-coding, long intergenic non-coding and antisense genes were kept. Finally, the matrices of counts were filtered to keep only the spots under the tissue of the corresponding image datasets.

## Regression formula notation

Regression formulæ use the following special arithmetic rules. The 1 denotes an intercept term. *a*:*b* denotes an interaction term for covariates *a* and *b*, and *a*\**b* expands to *a* + *a*:*b* + *b*. Addition and interaction are idempotent, *a* + *a* = *a*, and *a*:*a* = *a*.

## Spatial Transcriptome Decomposition

### Negative binomial decomposition

#### Core model

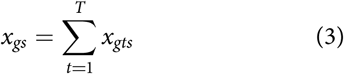

and that these in turn are negative binomially distributed,

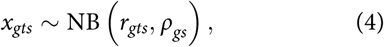

with rate and odds parameters *r*_*gts*_ and *ρ*_*gs*_, according to eq. (5),

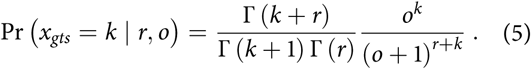

Notably, the odds parameters *ρ*_*gs*_ are restricted to not depend on the factor because from this follows a simplified likelihood, *x*_*gs*_ ∼ NB (Σ_*t*_ *r*_*gts*_, *ρ*_*gs*_); this allows us to marginalize over *x*_*gts*_ during inference. Furthermore, in practice, the odds parameters of our models are typically chosen to depend only on the gene, *ρ*_*gs*_=*ρ*_*g*_. This restriction is also present in other models, such as DESeq2, ZINB-WaVE, and scVI for which the dispersion parameters only depend on the gene [5, equation 1] [6, eq. 6][7, eqs. 4 and 5].

#### Rate and odds regression

In the framework, the logarithms of the rate and odds parameters are specified in terms of regression formulæ. The default regression formulæ are

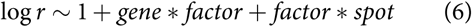

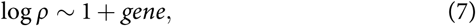

see the preceding section for an explanation of the formula notation. These regression formulæ correspond to the following regression equations:

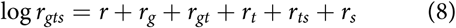

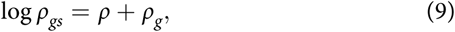

where the indices *g, t, s* denote covariate dependence on *gene, factor*, and *spot*, respectively.

#### Covariates

Aside from the above-mentioned covariates (intercept, genes, spots, and factors), it may often be necessary to include additional terms in the rate and odds regression formulæ. For example, when data of multiple sections are analyzed then *section*+*gene*:*section* terms can capture technical noise. Furthermore, by writing design files, the samples may be annotated with additional covariates according to the experimental design, and terms depending on these covariates may be used in the formulæ.

In this way, when sections from multiple different biological conditions are analyzed, for example different cancer types, then biological variation can be captured by *gene*:*cancer* terms. Thus, when data are available that control for different cancer types and that comprise multiple sections as repeat experiments, then the following rate regression formula may be appropriate:

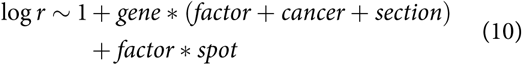

#### Probability distributions and hierarchies

In order to adapt the model to the experimental design of the specific application, the framework offers flexible modeling choices for the coefficient prior structure. Available prior distributions include the normal, beta, and gamma distributions; and arbitrary directed graphical probabilistic hierarchies can be built out of these.

#### Model specification

The graphical probabilistic structure of the model is specified using conventional mathematical notation in model specification files. The regression for the rate and odds parameters can be specified either directly in terms of equations or in terms of formulas that get translated into equations. The following is an example model specification file:

~~~
# Rate equation
rate = rate()+rate(gene)+rate(spot)+rate(gene, type)+rate(spot, type)+rate(type)
# Odds equation
odds = odds()+odds(gene)
# Coefficient distributions
rate()           ∼ Normal(0, 1)
rate(gene)       ∼ Normal(0, 1)
rate(gene, type) ∼ Normal(0, 1)
rate(spot)       ∼ Normal(0, 1)
rate(spot, type) ∼ Normal(0, 1)
rate(type)       ∼ Normal(0, 1)
odds()           ∼ Normal(0, 1)
odds(gene)       ∼ Normal(0, 1)
~~~

Note that type corresponds to the transcriptomic factor.

The regression equations may be arbitrary mathematical expressions composed of sums, differences, products, divisions, exponentiations, and logarithms.

Instead of the regression equations given above, the following formulas could be used equivalently:

~~~
# Rate formula
rate := 1+type*(gene+ spot)

# Odds formula
odds := 1+gene
~~~

Syntactically, expressions with an equality sign, =, such as in the first example, are parsed as equations, while expressions with := are parsed as formulas. Finally, expressions with a tilde symbol, ∼, are parsed as probability distribution specifications. If no probability distribution is specified for a coefficient, then it is assumed to be distributed according to the standard normal distribution.

#### Engineering model flexibility

Technically, our method’s modeling flexibility is enabled by representing the abstract syntax tree (AST) corresponding to the user-specified mathematical expressions in the regression equations and computing symbolic derivatives for gradient-based optimization. Function and derivative evaluation relies on just-in-time code generation and compilation for the expressions from the AST utilizing the LLVM compiler framework [8].

#### Optimization

The framework offers several parameter optimization schemes based on likelihood gradients, including RPROP [9], AdaGrad [10], and ADAM [11] optionally with a Nesterov-type momentum term [12, 13].

#### Stochastic gradient

Stochasticity is injected into the learning process by two means: we compute a stochastic approximation to the gradient by randomly ignoring counts *x*_*gs*_ during gradient calculation. Furthermore, it is often beneficial to dynamically down-sample the counts so that all experiments have the same read-per-spot ratio. The stochastic gradient approximation linearly speeds up the computation but more importantly helps avoid over-training. Down-sampling of counts to equate the read-per-spot ratios avoids uneven likelihood contributions across the experiments due to sequencing depth, which otherwise frequently results in factors focussed more towards explaining samples with higher sequencing depth.

### Staging

Optimization is done in multiple rounds, in which increasing numbers of parameters are included into the optimization. In the first stage only global, scalar coefficients are optimized. In the second stage, we additionally optimize scalar coefficients that depend on further covariates such as *section* or *individual*. From the third stage optimization includes gene-dependent and spot-dependent coefficients that do not depend on further covariates. Stage four also includes gene-dependent and spot-dependent coefficients that depend on further covariates. The fifth stage finally optimizes all coefficients, including gene- and factor-dependent ones, as well as spot- and factor-dependent ones.

Stages one to four perform fifty iterations of gradient updates, the fifth stage performs 2000 iterations.

### Visual summarization and clustering

#### Visual summarization

Spatial patterns across many features—regardless whether genes or factors—can be visually summarized with dimensionality reduction techniques from machine learning, see fig. S2. In this context, we consider matrices that have rows for every spot and columns for every feature (which could be genes or spatial factor activities). Utilizing t-distributed stochastic neighbor embedding (t-SNE) [14] or similar methods, such as principal component analysis (PCA) or uniform manifold approximation and projection (UMAP) [15], the number of columns of the matrices is reduced to three. The data are then rescaled into the unit cube and the rows used as coordinates in color space to colorize the spots in a spatial plot. When spots are colored in this way, similar colors indicate similar gene expression or similar factor activities, indicating similar cell type composition. We refer to such plots as maps of transcriptomic anatomy if the features reflect transcriptome-wide gene expression data.

#### Hierarchical clustering

Aside from visualizing cell type composition by dimensionality reduction of spatial activity maps, it is also possible to apply hierarchical clustering to the spatial activity maps. Unlike the quantitative visualization that dimensionality reduction based approaches yield, clustering partitions the spots into discrete sets, which can be useful for down-stream analyses.

### Differential gene expression analysis

Differential gene expression (DGE) analysis is performed as follows. The analyst decides on a suitable number of clusters by inspecting the hierarchical clustering results. For the chosen number of clusters, the spots are then partitioned into sets for each cluster. For all pairs of clusters, a pair-wise DGE analysis is performed using DESeq2 [5].

The performance of the scaling factor determination of DESeq2 deteriorates with increasing number of spots because it can only utilize genes that have non-zero counts throughout all spots, and the probability of observing no zero count for a given gene in each of *n* samples decreases exponentially in *n*. Therefore, instead of using all spots of a given cluster (which can number in the hundreds or thousands), we randomly select subsets of about 50–100 spots. This is justified based on the assumption that the spots within a cluster are interchangeable.

### Functional enrichment analysis

To perform functional enrichment analyses, we make use of the R package enrichR, which provides an R interface to the web-based functional enrichment analysis tool Enrichr [16]. The following ontologies were included:

- GO Molecular Function 2017
- GO Cellular Component 2017
- GO Biological Process 2017
- KEGG 2016
- Reactome 2016
- Allen Brain Atlas up

## Design and analysis of biological data

The experimental design for the olfactory bulb and the brain samples is given in table T1 and table T2. For the olfactory bulb samples, we discard the sex covariate since it is confounded with individual B. We thus use the following rate regression formula for both the olfactory bulb and brain analyses:

**Table T2:** Design of mouse coronal brain experiments.

| Section | Individual | Sex | Reference |
| --- | --- | --- | --- |
| 1 | A | ♂ | here |
| 2 | A | ♂ | here |
| 3 | B | ♂ | here |
| 4 | B | ♂ | here |

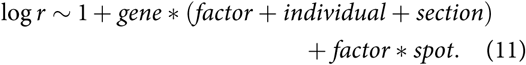

Analyses were performed with version 0.3-111-g40dcf70 of std-nxt and the following command:

~~~
std-nxt --adjdepth --stage 50 \
 --minread_spot 100 --dropout 0.2 \
 -v -i 2000 -r 10 -t 20 --optim adam \
 --design design.txt --model model.txt
~~~

## Synthetic data experiments

### Model

The synthetic data is generated according to the following model:

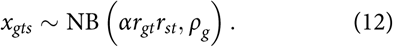

For notational brevity, we will use *φ*_*gt*_ and *θ*_*st*_ in place of *r*_*gt*_ and *r*_*st*_ in this section and denote them in matrix form as Φ and Θ, respectively.

### Measuring performance

We evaluate the success of the decomposition by computing the Pearson correlation between the expected number of reads generated by the true and inferred models in the gene-factor and spot-factor marginals. That is, we use 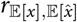 as a performance measure, where either

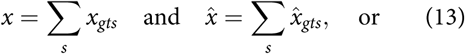

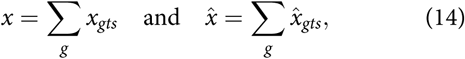

and where *x*_*gts*_ and 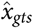 are the number of reads generated by the true and inferred models, respectively, for a given gene *g*, transcriptomic factor *t*, and spot *s*.

### Ground truth parameters

In each experiment, we generate data for |*T*| =10 factors and let

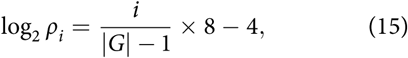

where *i* ∈ [0, 1 … |*G*| −1] and |*G*| = 1000.

We examine model performance along four dimensions: the average number of reads per spot, the number of spots, the heterogeneity of gene profiles, and the heterogeneity of activity maps. In order to measure the latter two quantities, we consider the random vector (*G, T, S*), defined so that *P*(*G*=*g, T*=*t, S*=*s*) is equal to the relative frequency of reads for a given gene *g*, factor *t*, and spot *s*:

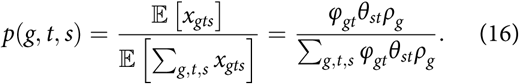

We define the heterogeneity of the gene profiles and spatial activity maps, respectively, as

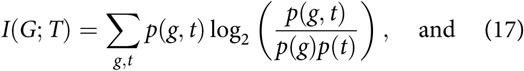

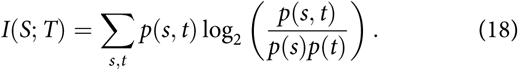

#### Sampling

We use the following procedure to sample the parameters *α*, Φ, and Θ for an experiment with an average of *n* expected reads per spot, |*S*| spots, and factor heterogeneities *i*(*G*; *T*) and *i*(*S*; *T*):

1. Initialize Φ to a |*G*|×|*T*| matrix and Θ to an |*S*|×|*T*| matrix with each entry set to 1/|*T*|.
2. Sample Φ and Θ using the Metropolis-Hastings algorithm on the density *p*(Φ, Θ) ∝ *e*^−*E*^, where

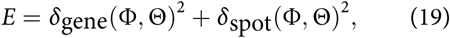

and

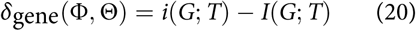

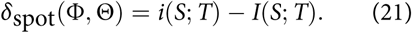 In order to improve convergence speed, we tune the proposals based on the values of (20) and (21). Formally, we use a time dependent proposal function *N*_*t*_(Φ, Θ, *S*_*t*_), where

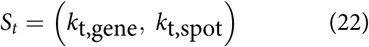

denotes its state at time *t*. We let its initial state be *S*_0_ = (0,0). *N*_*t*_ generates new proposals and modifies its state according to the following procedure:

a. Choose one of

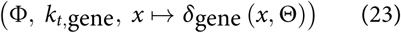

and

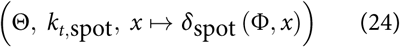

at random with equal probability. Call the chosen triplet (*M, k, f*).
b. Select a random set of *m* rows in *M*, where *m* = 0.01× rows (*M*)?. Each row is drawn equiprobably from the set of all rows in *M* without replacement. Define 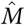 to be *M* with the selected rows replaced with *m* samples from a Dir (*e*^*k*^ *u*) distribution, where *u* is a |*T*|-dimensional vector of ones.
c. Let 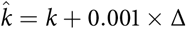, where

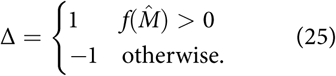
d. The new proposal is

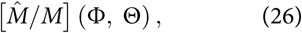

where the operation [*x*/*y*] *z* denotes replacing variable *y* with *x* in expression *z*.
e. The new state of the proposal function is

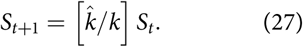
3. Given Φ and Θ, we can solve for the scaling parameter *α*:

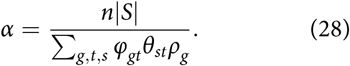

### Inference

Using the data generated by the model in the preceding section, we infer the parameters of the model

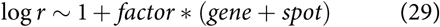

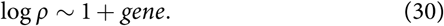

We run `std-nxt` with the following command:

~~~
std-nxt -t10 -i2000 -m model.txt \
 --optim=adam --adam_nesterov counts.tsv
~~~

#### Comparison to Poisson regression

We run the method of Berglund *et al.* [17] with the following command:

~~~
std -t10 -i2000 counts.tsv --sample=\
contributions,phi,phi_prior,phi_local,\
theta,theta_prior,spot,baseline
~~~

### Poisson model

To better understand how our method compares to the one of Berglund *et al.* [17], we substitute eq. (12) to conform to their inference model:

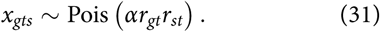

Sampling of *α, r*_*gt*_, and *r*_*st*_ and inference are performed analogously to what has been described above. The results of the performance evaluation are shown in fig. S11.

## Comparison with related methods

We assessed the performance of two related methods, scVI [7] and ZINB-WaVE [6], on the same datasets analyzed in this manuscript.

### Front-end code

Both methods provide library routines, implemented in Python and R, respectively, for which the user has to write front-end code. The front-end code we wrote for scVI is available at https://github.com/maaskola/run_scVI and for ZINB-WaVE at https://github.com/maaskola/ run_ZINB-WaVE.

### Patching scVI

In addition to writing front-end code, we also had to patch the scVI code base slightly to enable two features not present in the upstream code:

1. to operate on local count matrix files, and
2. to pass the spot coordinate information through dataset concatenation operations.

Our patched version of scVI is available at https://github.com/maaskola/scVI.

### Covariate usage

ZINB-WaVE was supplied with exactly the same covariate information (covariates for section and individual) as our own method, STD. Only one label covariate can be used in the case of scVI, and so we decided to supply scVI with the section covariate for each spot. The same number of 20 factors, or hidden dimensions, was used for scVI and ZINB-WaVE as in our STD analyses.

### Additive and multiplicative factor analysis

We next explain the difference between additive (STD) and multiplicative (scVI and ZINB-WaVE) factor analysis. In additive factor analysis, the factors’ contributions add up to yield the observations. Our manuscript’s equation 1 reflects this immediately.

In multiplicative factor analysis, the various factors do not add up in a similar way. Rather, the factors simultaneously act to increase or decrease the expected counts in a multiplicative fashion.

Thus, the additive factors of STD yield an interpretation of the results in terms of the relative frequencies of reads in a spot explained by a given factor. These are numbers between 0 and 1, where 0 indicates that no reads are explained by the factor, and 1 indicates that all reads are explained by the factor. On the other hand the multiplicative factors of scVI and ZINB-WaVE result in coefficients which are real numbers—i.e. they can be negative—and thus a given factor can act positively or negatively on the expression pattern in a given spot. Additive factors in STD thus express to which extent a given factor is present in a given spot, while multiplicative factors in scVI and ZINB-WaVE express whether a given factor acts positively or negatively in a given spot.

Notably, additive and multiplicative factor analysis are both efficiently computable using simple linear algebra. In the case of additive factor analysis, the matrix multiplications are taken after exponentiating the logarithmic expression of the emission parameters, while for multiplicative factor analysis the matrix multiplications are taken inside the logarithmic expressions for the emission parameters.

Both are in principle valid approaches to modeling the data. We posit here on theoretical grounds that due to ST expression measurements having the character of mini-bulk samples (since they receive mRNA from multiple cells) that additive factor models are more appropriate to model ST count data. Below, we additionally provide empirical evidence for the superiority of modeling ST data with additive factor analysis.

Usage of additive and multiplicative factors is not mutually exclusive, and both kinds of factors can be used within one model.

### Visualizing latent spaces of scVI and ZINB-WaVE

Indexed by the spot and factor dimensions, the latent space representations of scVI and ZINB-WaVE are spatial objects. We can thus visualize them in a way similar to the STD results, with a slight difference. For STD there exists an interpretation in terms of contributions due to the different factors that add up to explain the observations. But since components of the scVI and ZINB-WaVE latent space representation can be negative, no such interpretation exists for them. For this reason it is not possible to visualize scVI and ZINB-WaVE results by displaying for each spot the relative frequencies of the factors’ contributions, as we can do for STD. Instead, we can directly visualize the (logarithmic) latent space representation. Regardless of interpretational differences between additive and multiplicative factor analysis, we can summarize the latent space representations using t-SNE or related dimensionality reduction techniques as we do for the STD spatial factor activities.

### Results

Figures S14b and S16b visualize the scVI latent space representations of the olfactory bulb and brain samples. The corresponding ZINB-WaVE results are in figs. S15b and S17b. t-SNE summarizations of the latent spaces are shown in figs. S14c and S16c and figs. S15c and S17c. UMAP summarizations of the latent spaces are shown in figs. S14d and S16d and figs. S15d and S17d. Hierarchical clustering results for the latent spaces are shown in figs. S14e and S16e and figs. S15e and S17e.

Considering the t-SNE and UMAP summarizations first, we find that analysis results of scVI and ZINB-WaVE are useful, but exhibit stronger residual batch effects than when using STD. Similarly, hierarchical clustering of the latent space representations to the same number of clusters as with the STD spatial activities yields less consistent results across the samples when compared with STD.

The STD spatial activity maps presented in the main manuscript reveal that a given factor can only be absent or enriched in a region. Contrarily, with multiplicative factor analysis, factors can also act negatively. This is noticeable in the scVI and ZINB-WaVE results by examining the visualization of the latent space representations closer. Most factors represent an axis in which one or multiple tissue components form one end of the axis, and another or multiple other tissue components form the other extreme. For example, factor 1 in the ZINBWaVE results on the brain sections corresponds to the axis hippocampal field versus white matter; factor 2 corresponds to the axis of thalamus versus cortex. Some of the factors represent rather complex combinations, such as factor 15 of the same analysis, which has the reticular nucleus and the dentate gyrus at one end of the spectrum and the hypothalamus and cortical layer 6 at the other end. It is thus obvious that multiplicative factors need to be combined in the right amounts so that unwanted positive and negative contributions cancel.

As in our STD analysis results, we find scVI and ZINB-WaVE to yield higher entropy in samples three and four of the brain dataset. Visually, it appears that this unwanted effect is stronger in the scVI and ZINB-WaVE results. Overall, the scVI results appear slightly noisier than those of ZINB-WaVE or STD.

---

**Figure S1:**
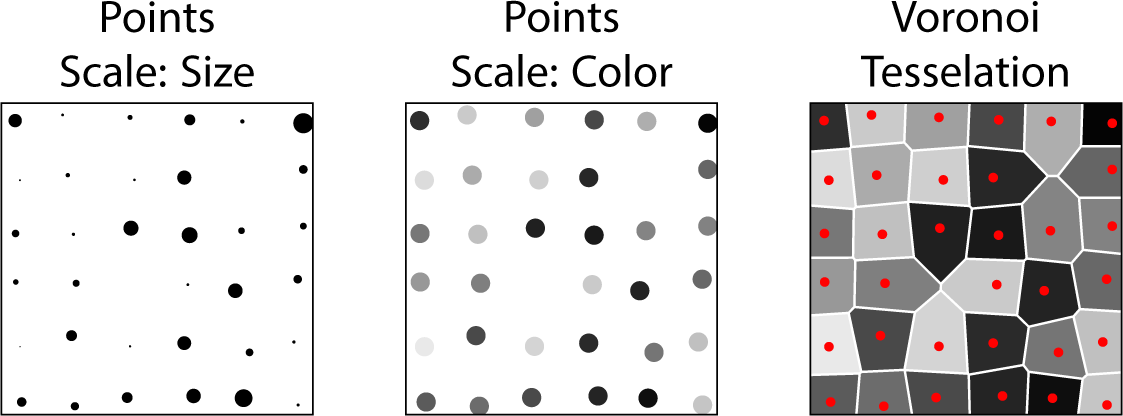
Three ways of visualizing spatial expression patterns. The geometries of the ST microarrays exhibit slight irregularities due to technical variability in the printing process: individual spots may be misprinted, and lack surface probes to capture mRNA; other spots may fuse with neighboring spots due to electrostatic attraction of the printing ink, and may have to be discarded. Points and vertices represent measurement points, with their plot positions corresponding to the spatial coordinates. Single channel information such as the expression of individual genes can be visually encoded in terms of size or color from a gradient palette. Alternatively, the Voronoi tessellation can be colored to convey the desired information.

**Figure S2:**
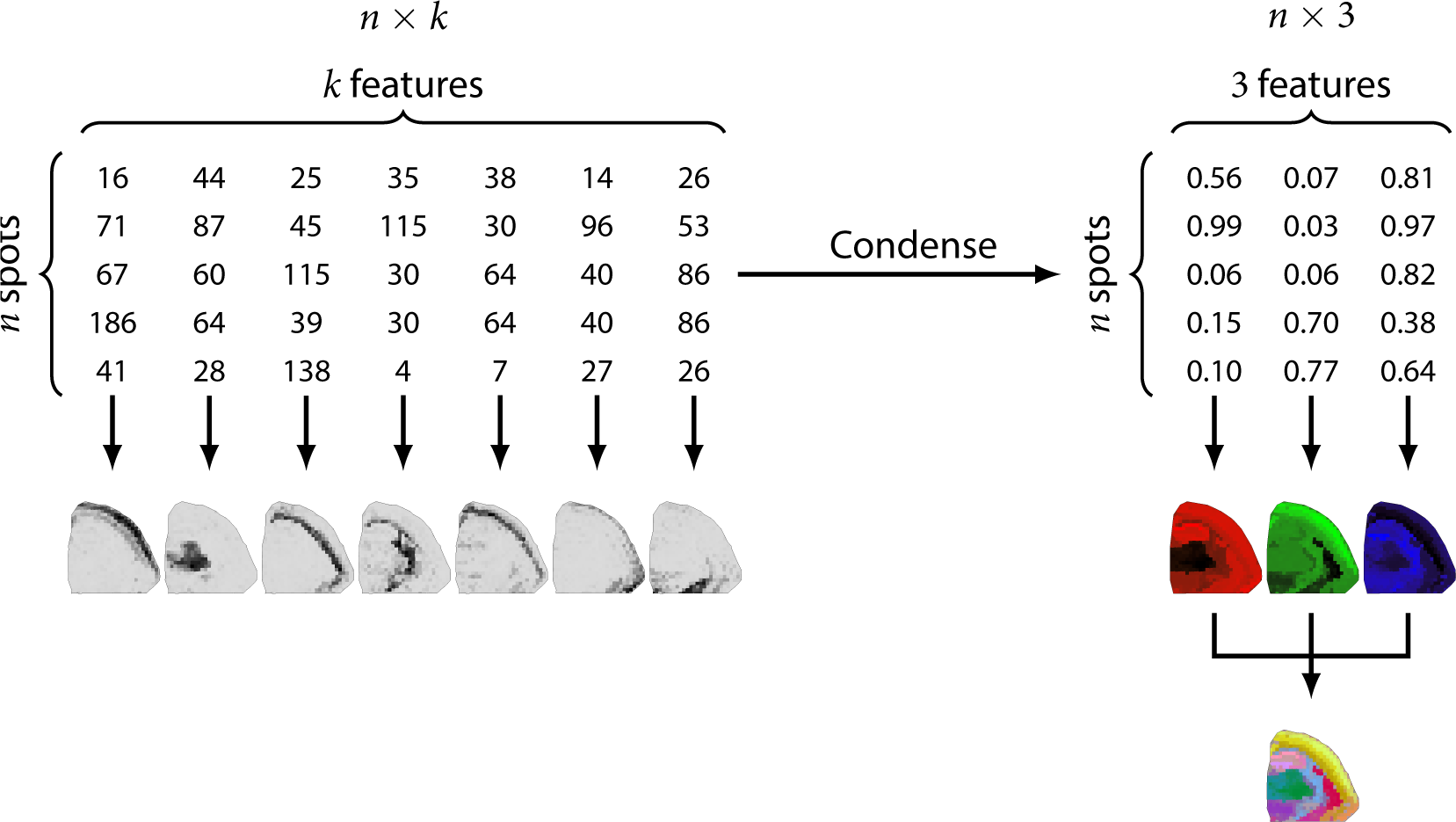
Visualizing complex spatial information by dimensionality reduction. The input matrices are *n* × *k* matrices corresponding to *n* spots and *k* features. The *k* input features may represent genes or factors, and the matrices typically contain sequencing read counts or factor activities. These *k* features may be visually represented with *k* single-channel images. Alternatively, the features may be combined and jointly visualized, making use of multiple color channels. By applying dimensionality reduction methods such as PCA, t-SNE, or UMAP, the *n* × *k* matrices are reduced to *n* × 3 matrices. The matrices are then rescaled to fit into the unit cube, and the three components are used as coordinates in color space to colorize spots in spatial plots.

**Figure S3:**
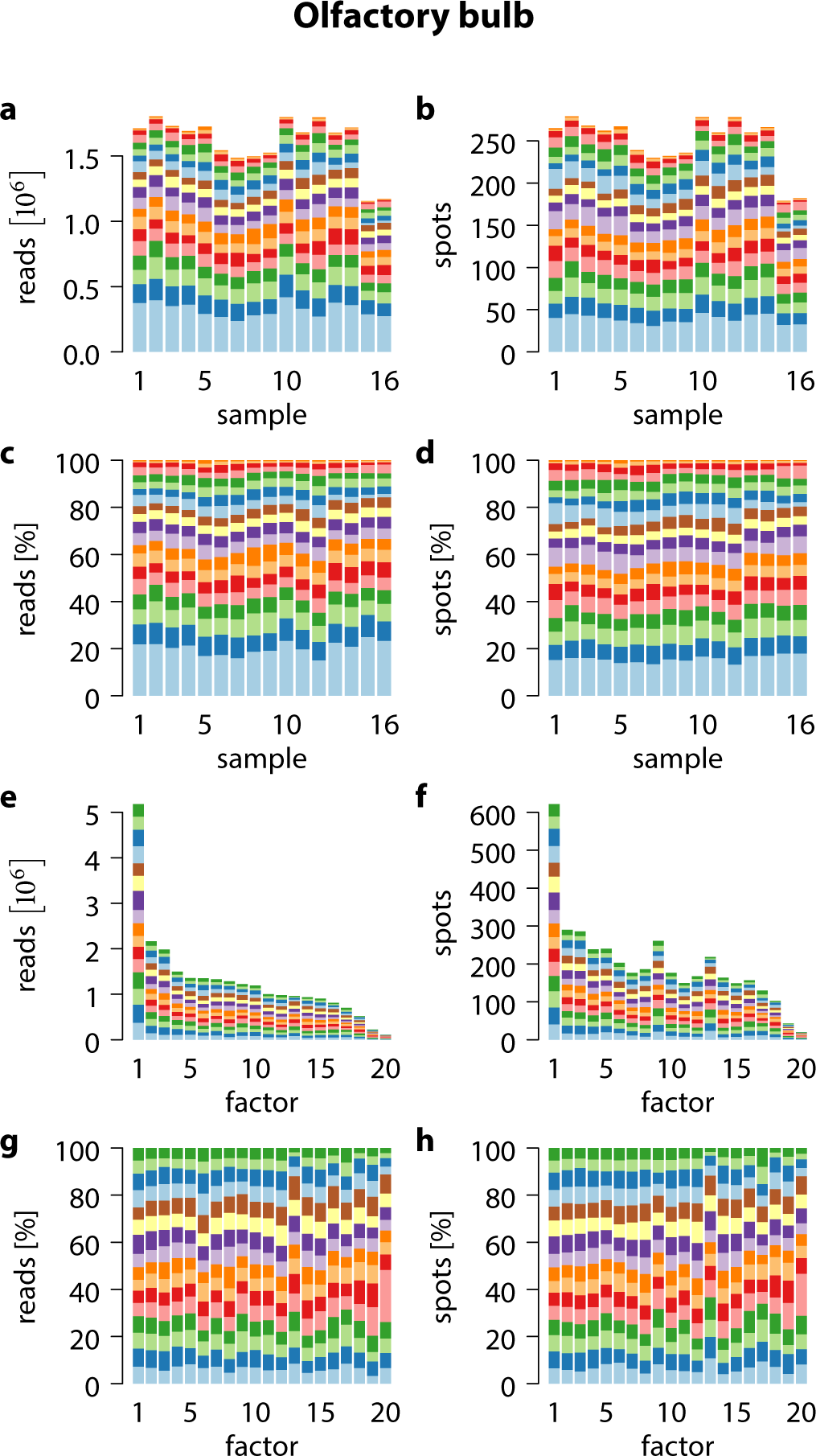
Reads and spots explained by factors across the mouse olfactory bulb samples. Colors denote factors (**a**–**d**) or samples (**e**–**h**).

**Figure S4:**
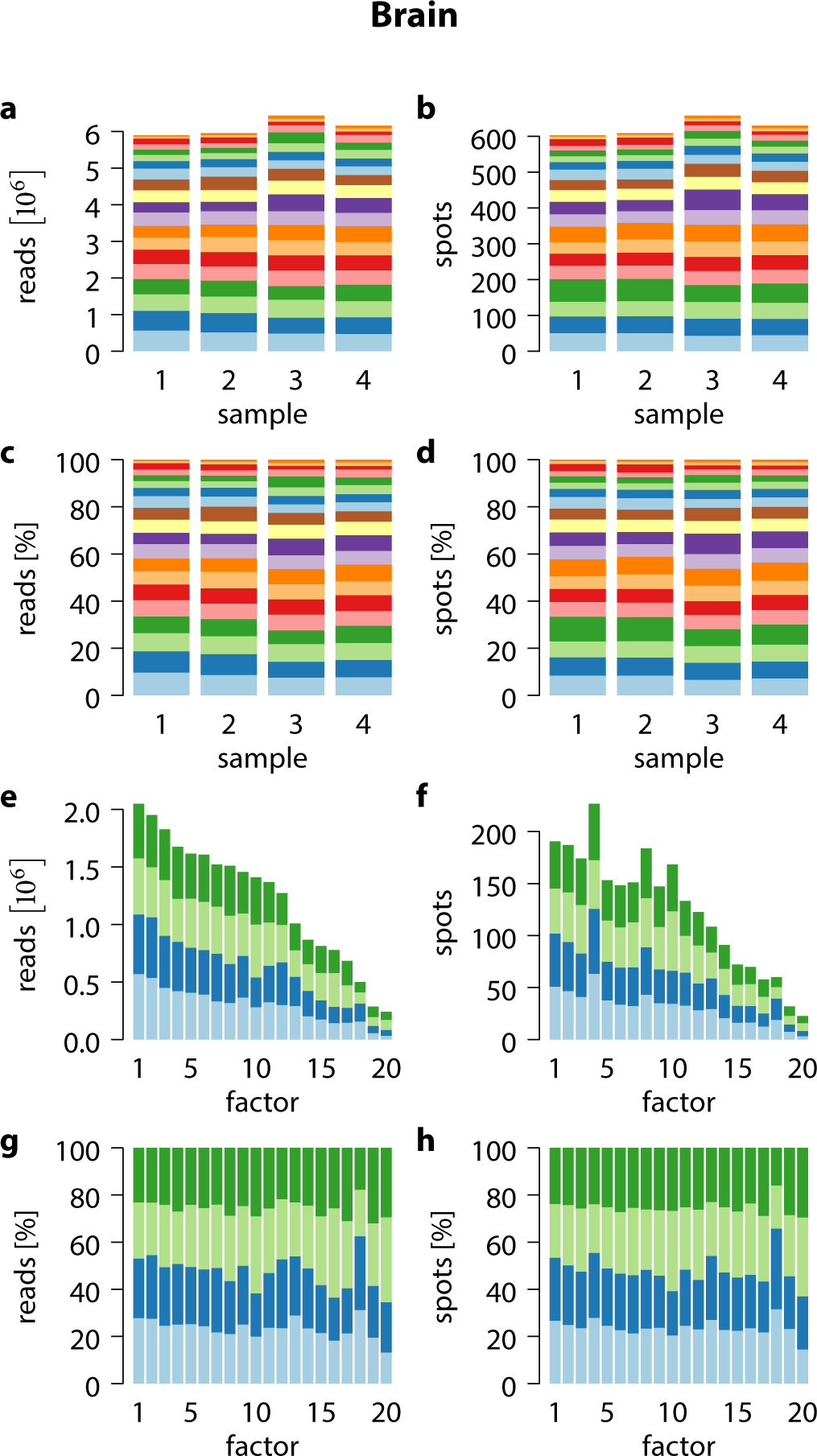
Reads and spots explained by factors across the mouse brain samples. Colors denote factors (**a**–**d**) or samples (**e**–**h**).

**Figure S5:**
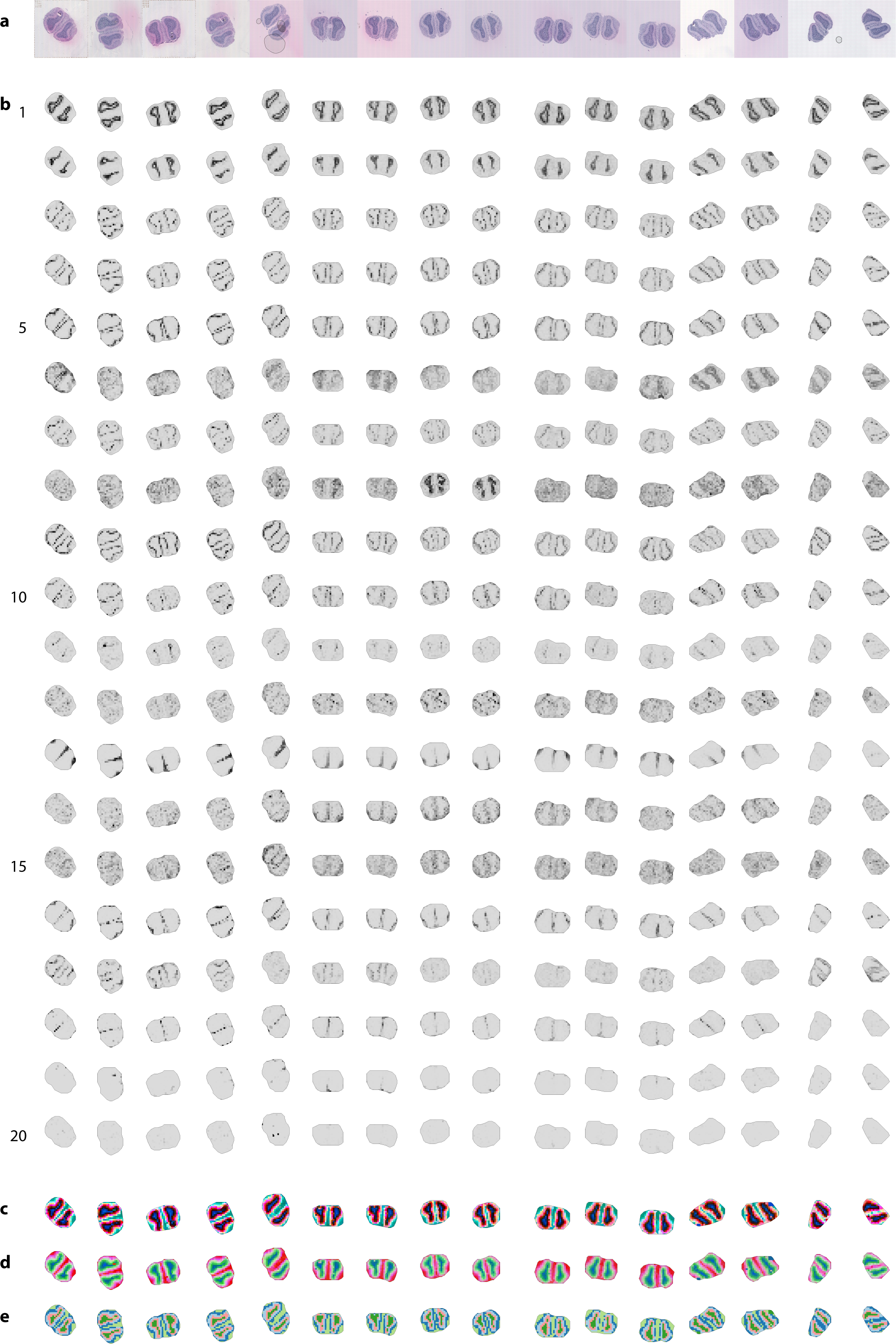
Transcriptomic patterns in mouse olfactory bulb sections. (**a**) H&E-stained microscopy images. (**b**) Spatial factor activity maps. (**c**) t-SNE summarization of factor activities. (**d**) UMAP summarization of factor activities. (**e**) Hierarchical clustering of factor activities into five clusters.

**Figure S6:**
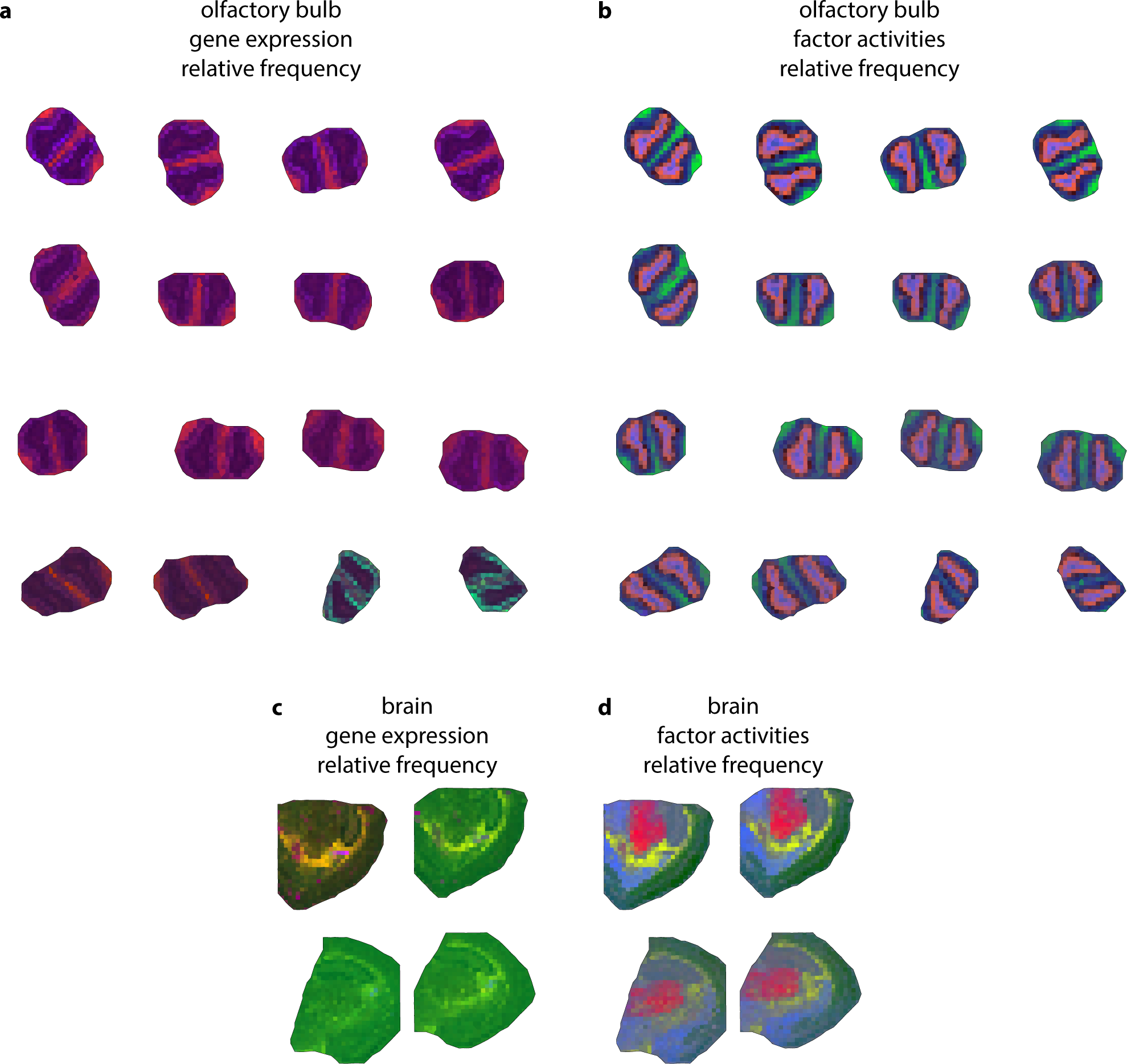
Summarization by principal component analysis (PCA) of gene expression (**a, c**) and factor activities (**b, d**) in mouse olfactory bulb (**a, b**) and mouse brain (**c, d**).

**Figure S7:**
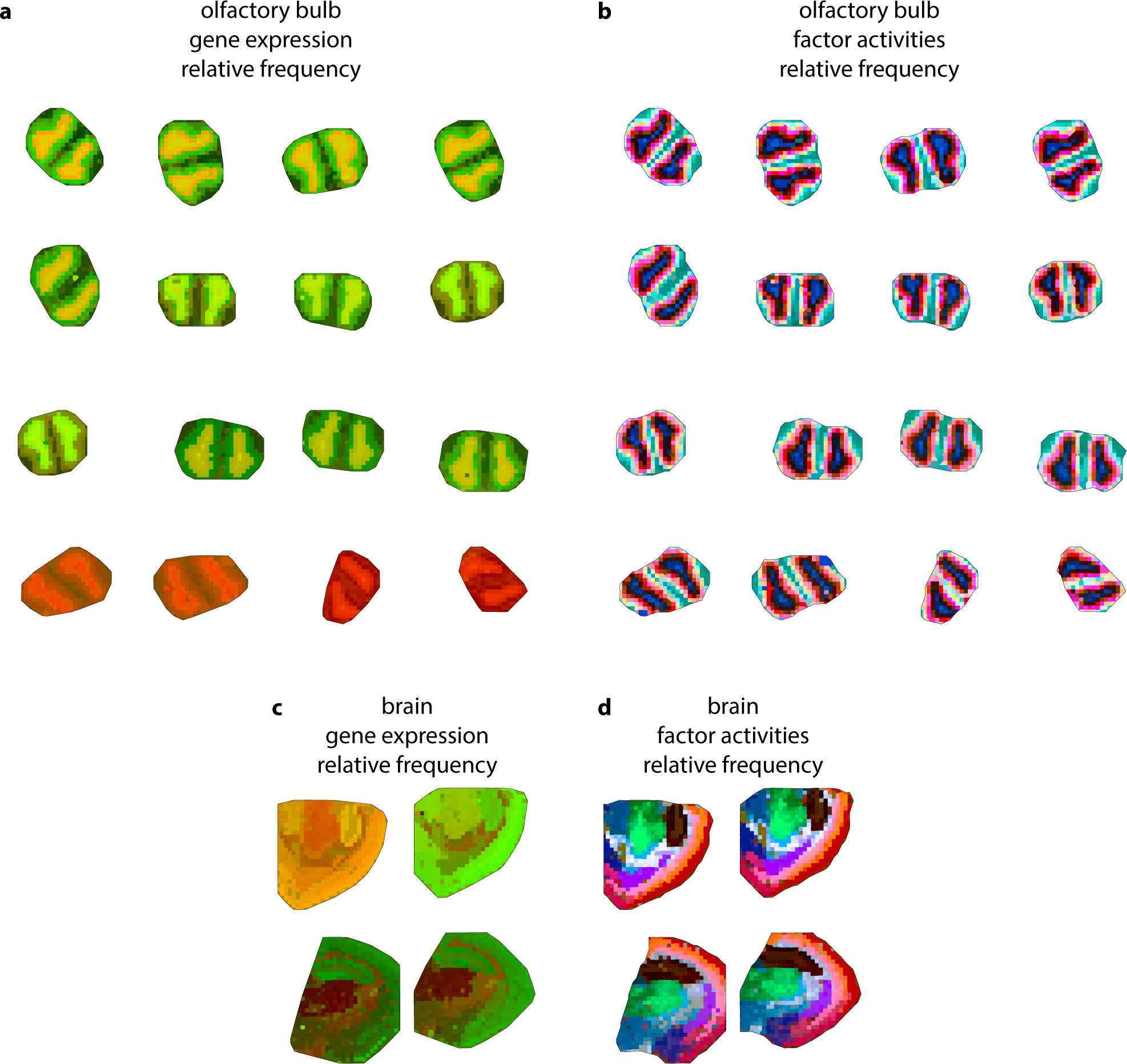
Summarization by t-distributed stochastic neighbor embedding (t-SNE) of gene expression (**a, c**) and factor activities (**b, d**) in mouse olfactory bulb (**a, b**) and mouse brain (**c, d**).

**Figure S8:**
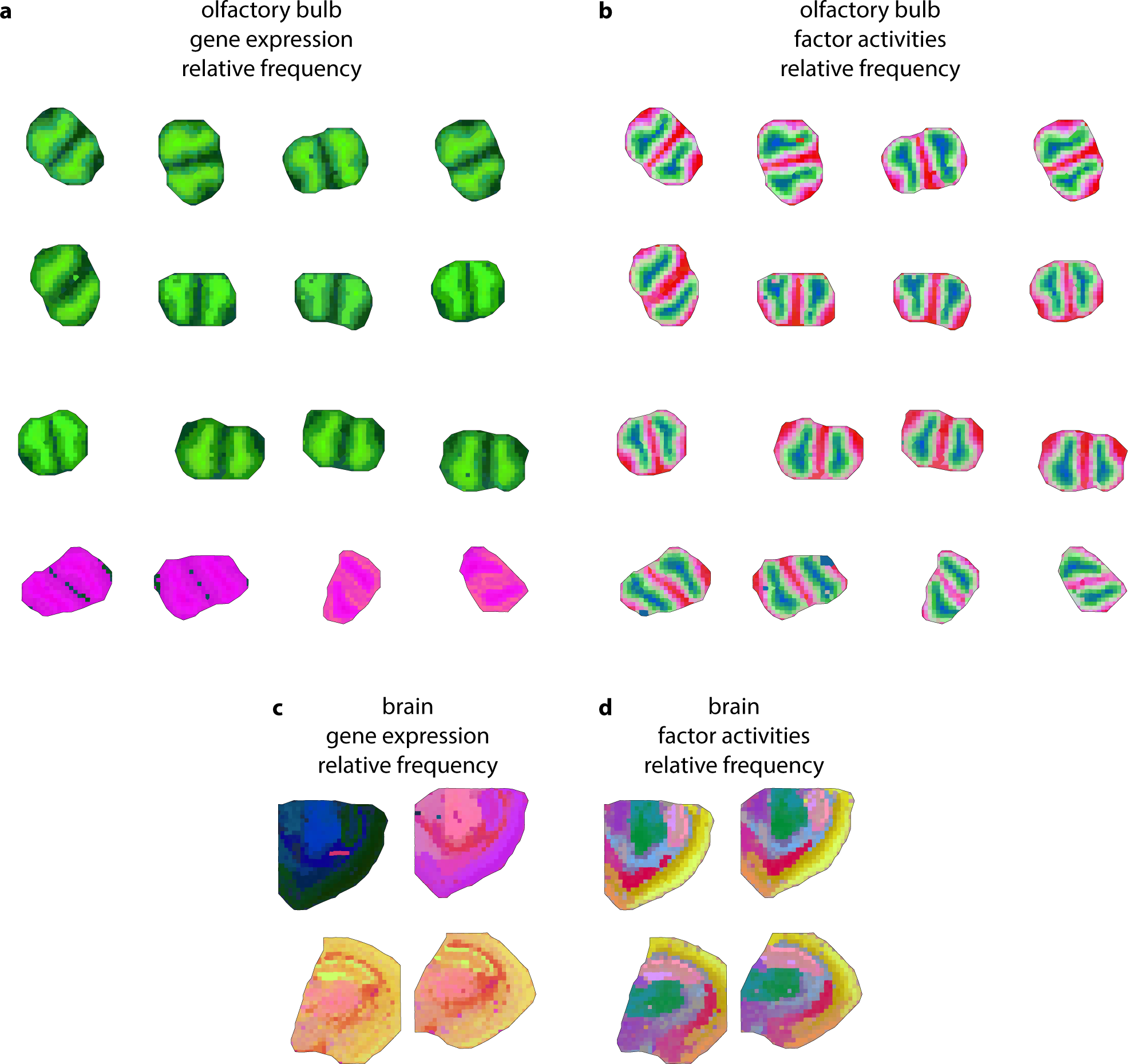
Summarization by uniform manifold approximation and projection (UMAP) of gene expression (**a, c**) and factor activities (**b, d**) in mouse olfactory bulb (**a, b**) and mouse brain (**c, d**).

**Figure S9:**
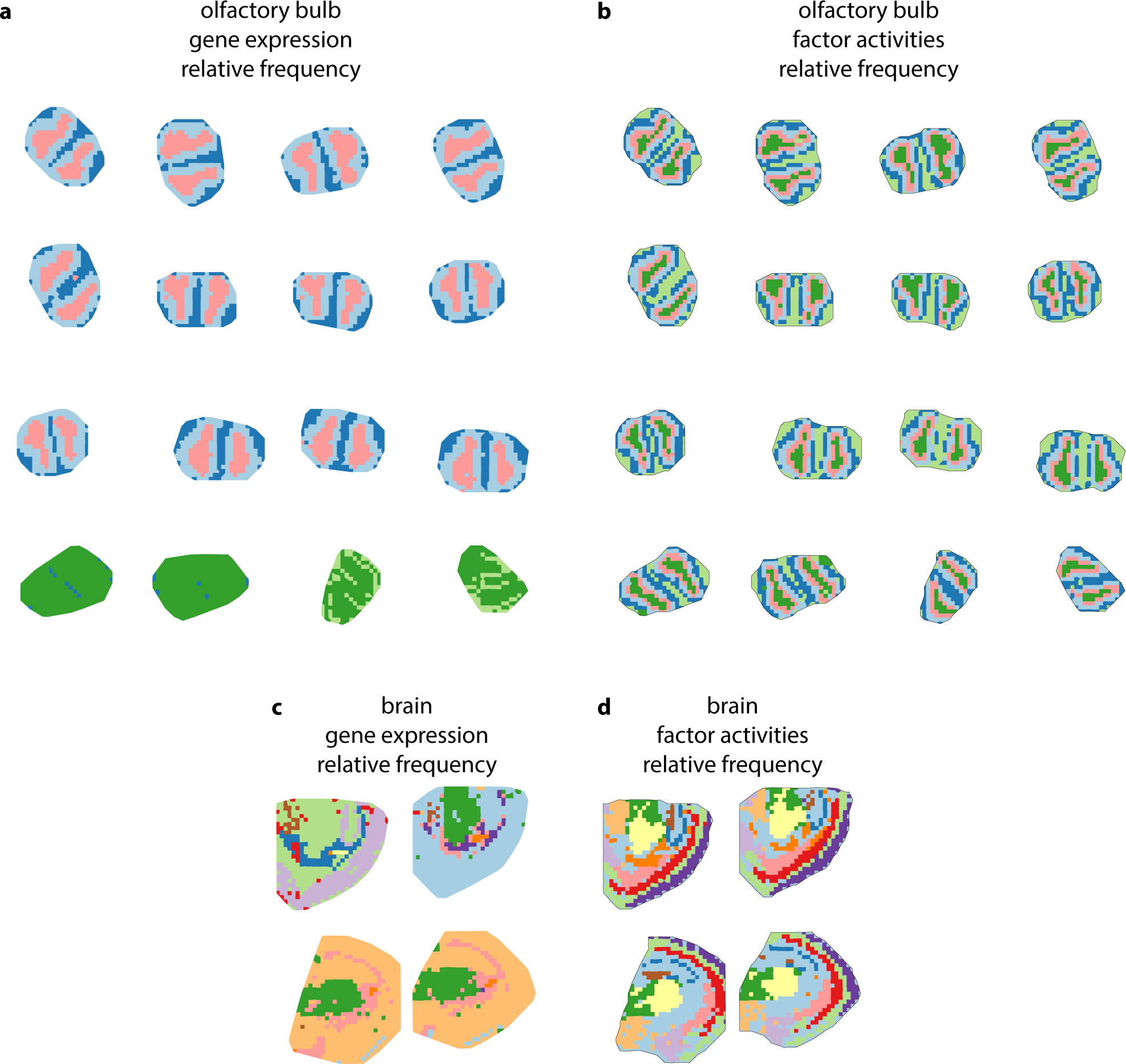
Hierarchical clustering of gene expression (**a, c**) and factor activities (**b, d**). Five clusters shown for mouse olfactory bulb (**a, b**) and twelve clusters for mouse brain (**c, d**).

**Figure S10:**
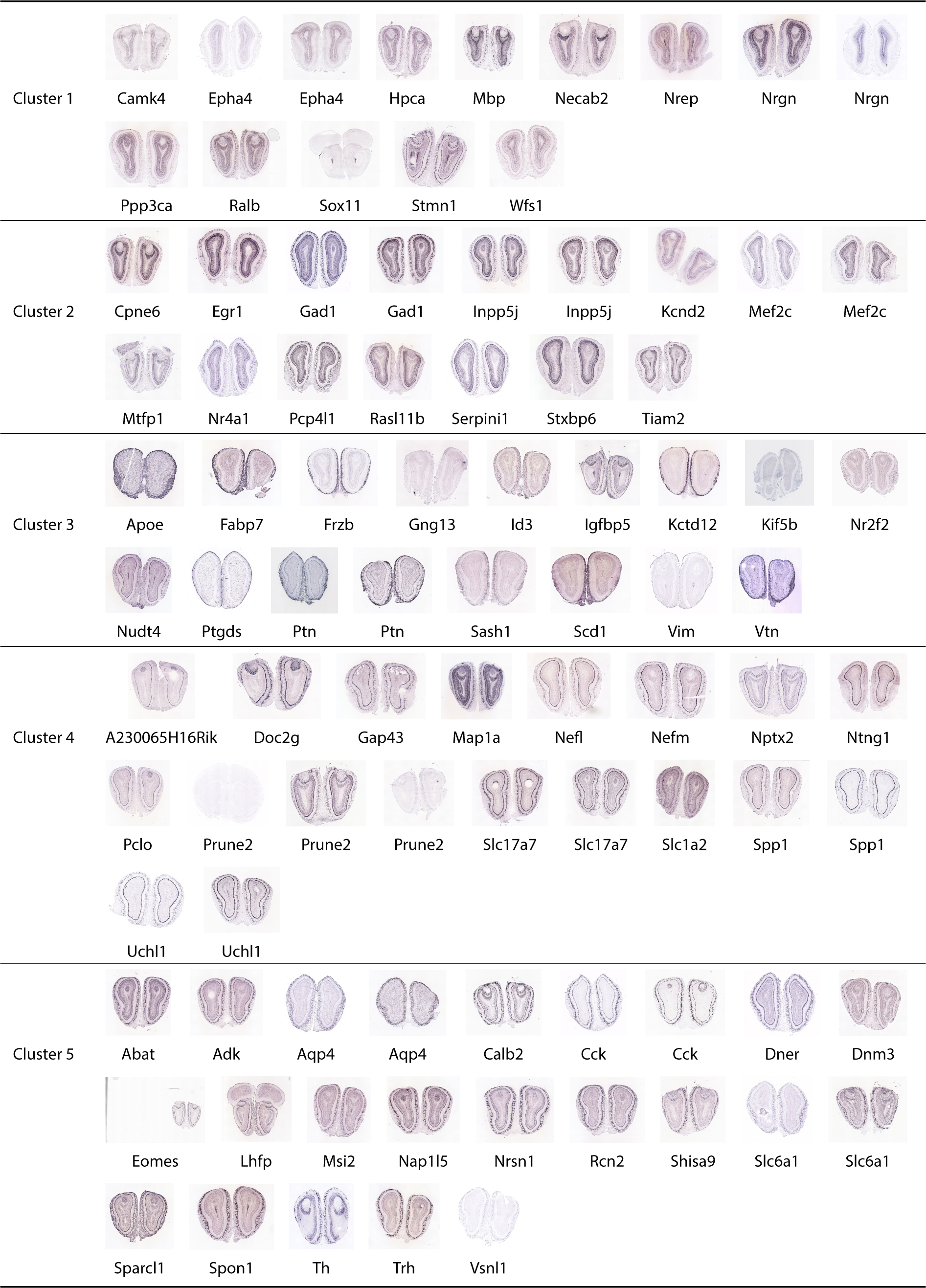
Spatial cluster specific genes in the Allen brain atlas [2]. Images from the Allen brain atlas for genes that are expressed significantly higher in one cluster versus all others. Based on clustering results shown in figs. S5e and S9b. From outside inwards, the clusters correspond to the olfactory nerve layer (layer 3), the glomerular layer (layer 5), the plexiform and mitral cell layers (layer 4), as well as two for the granular cell layer: a peripheral (layer 2) and a central one (layer 1).

**Figure S11:**
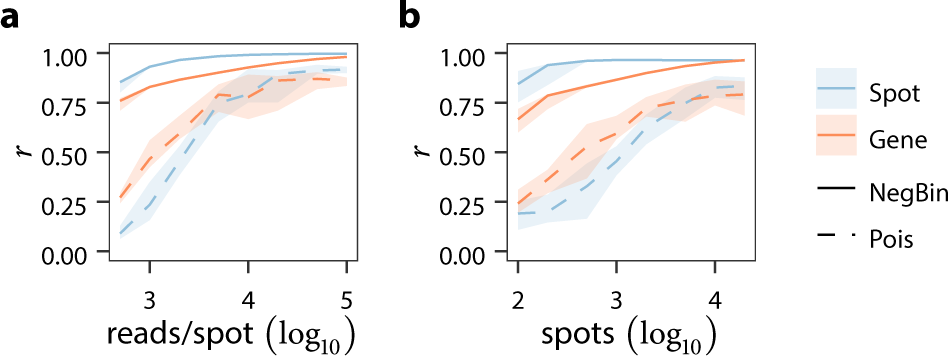
Performance evaluation on synthetic data using a Poisson source model. Plots show the correlation between the expected number of reads in the true and inferred models over the spot-factor and gene-factor marginals as a function of the number of reads per spot (**a**) and the number of spots (**b**) in the input data. Lines correspond to medians and shaded areas to minima and maxima. Solid lines show results for negative binomial decomposition, as described in this paper, while dashed lines show results for the Poisson regression framework of Berglund et al. [1].

**Figure S12:**
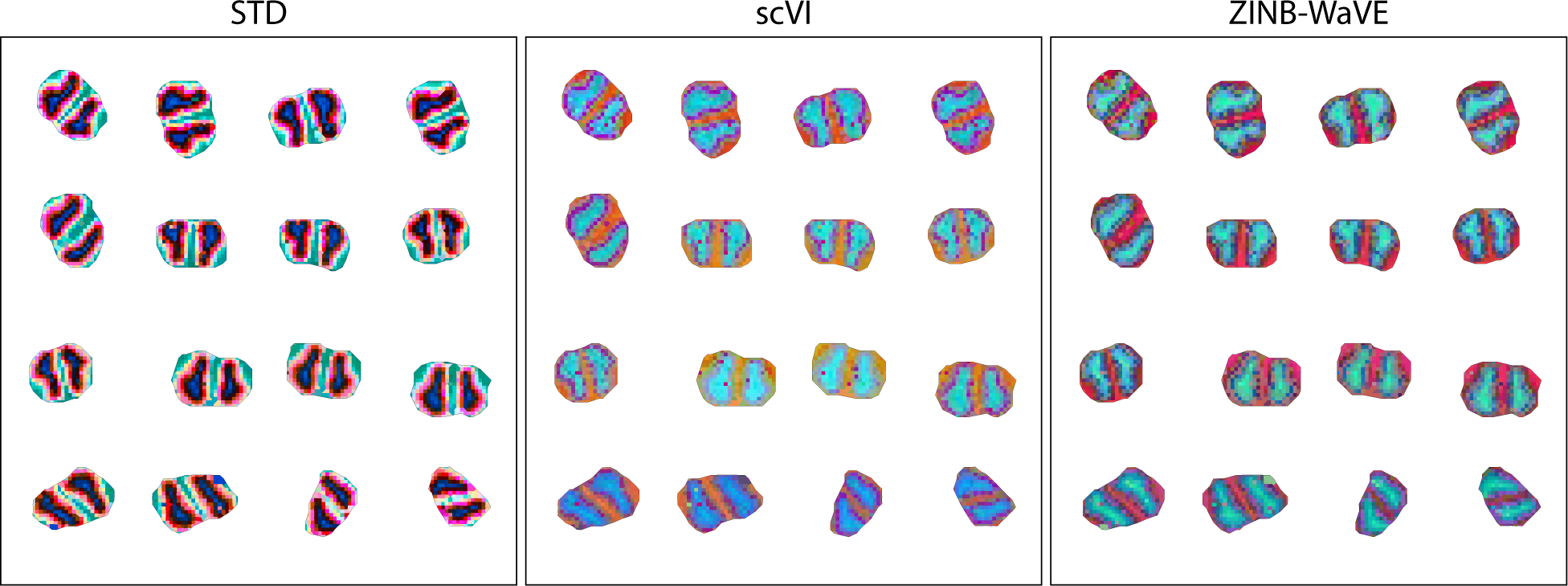
Analysis results of related methods for the mouse olfactory bulb sections, summarized using t-SNE. Analyses were performed with twenty factors, or twenty hidden dimensions, in each method.

**Figure S13:**
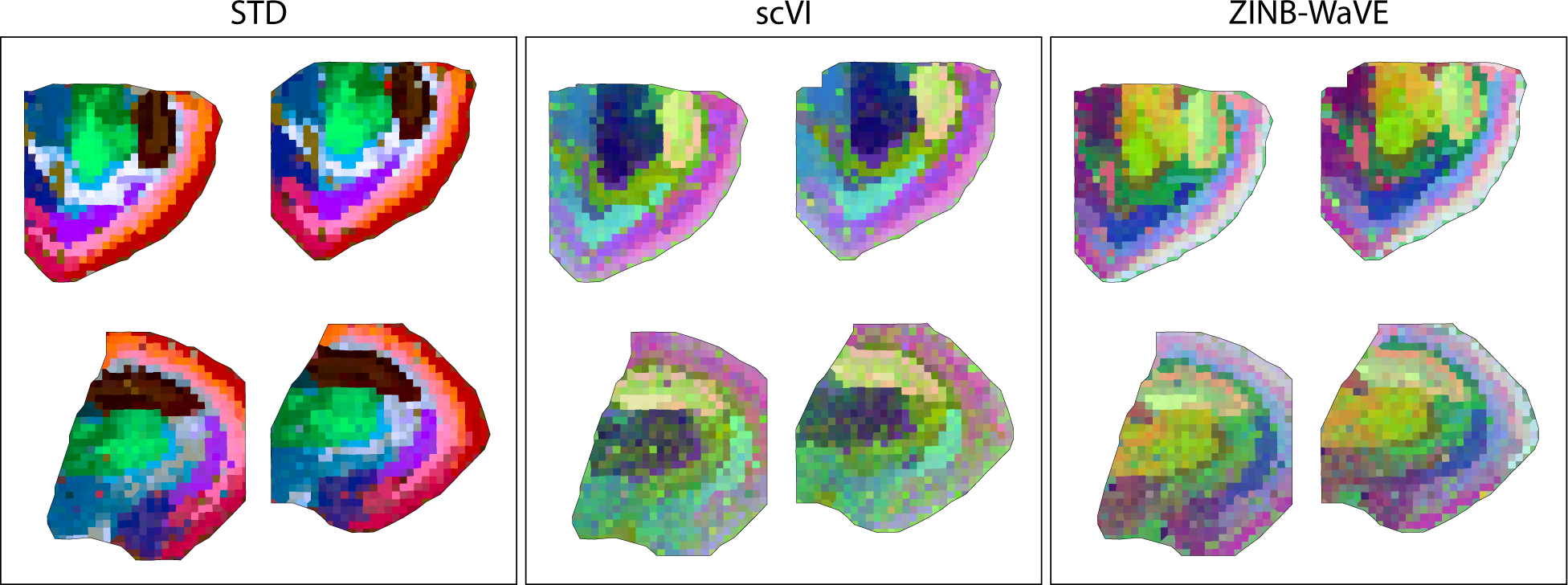
Analysis results of related methods for the mouse brain sections, summarized using t-SNE. Analyses were performed with twenty factors, or twenty hidden dimensions, in each method.

**Figure S14:**
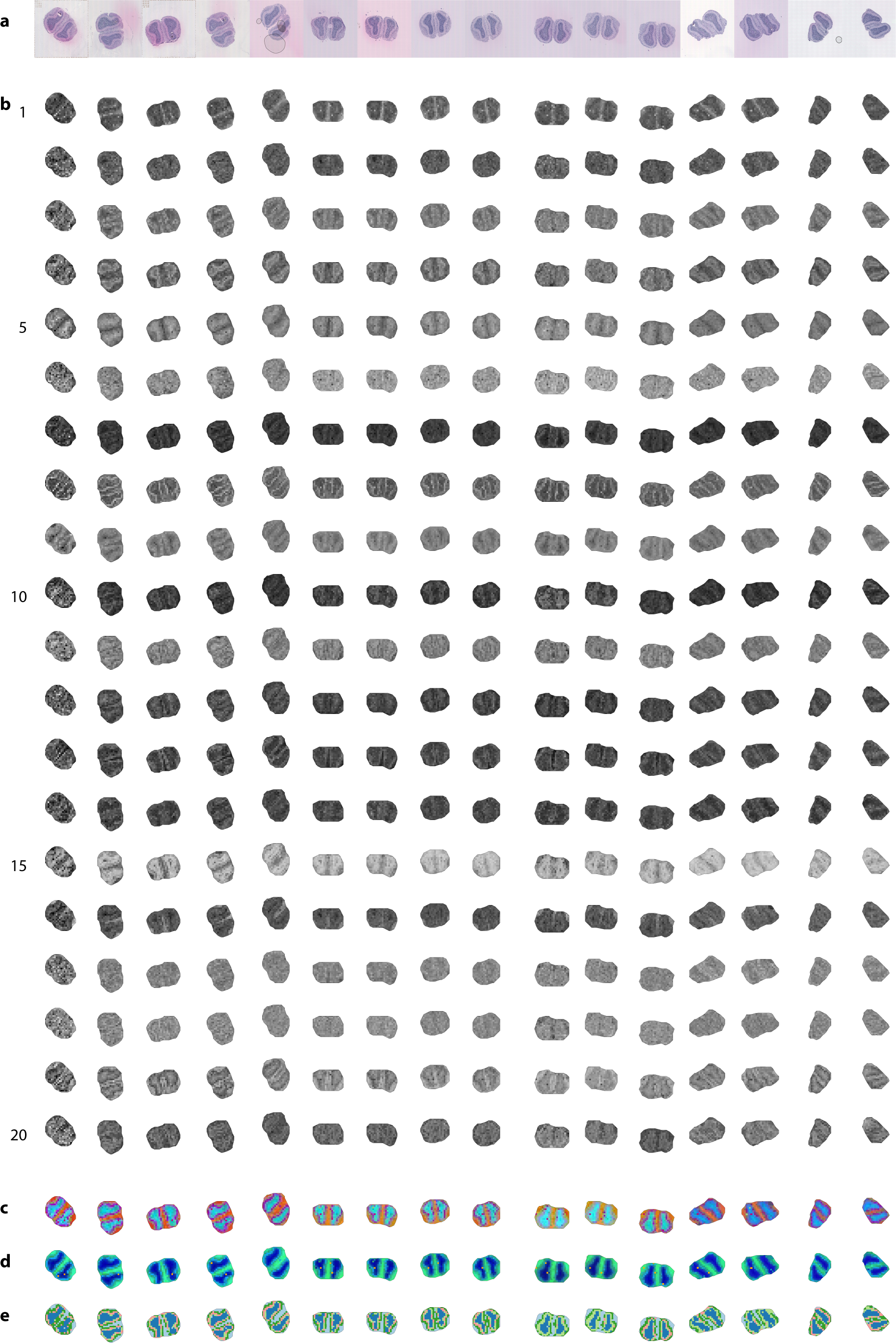
Analysis results for scVI in mouse olfactory bulb sections. (**a**) H&E-stained microscopy images. (**b**) Latent space representation of scVI. Note that the scVI latent dimensions are not sorted. (**c**) t-SNE summarization of scVI latent space. (**d**) UMAP summarization of scVI latent space. (**e**) Hierarchical clustering of scVI latent space into five clusters.

**Figure S15:**
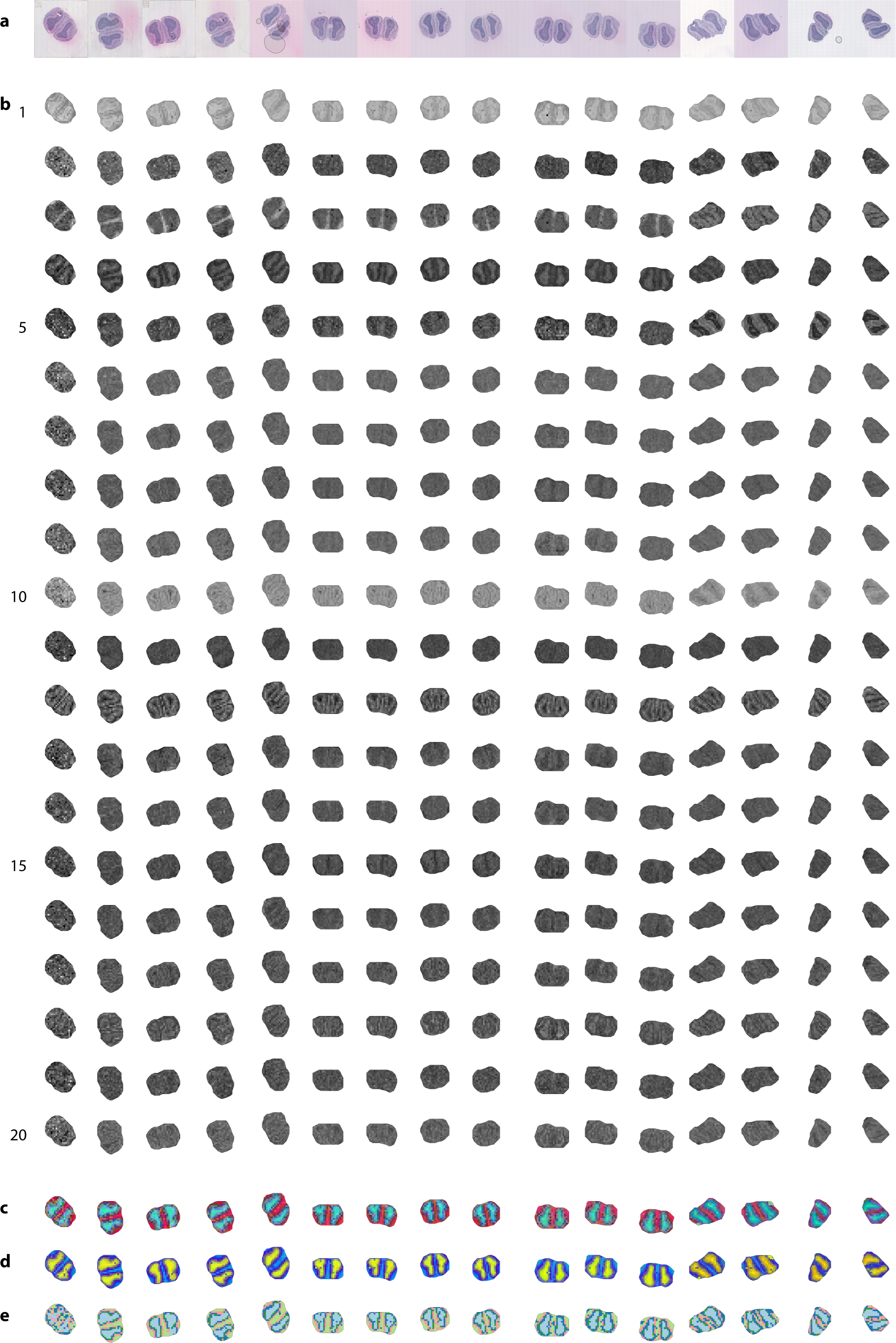
Analysis results for ZINB-WaVE in mouse olfactory bulb sections. (**a**) H&E-stained microscopy images. (**b**) Latent space representation of ZINB-WaVE. (**c**) t-SNE summarization of ZINB-WaVE latent space. (**d**) UMAP summarization of ZINB-WaVE latent space. (**e**) Hierarchical clustering of ZINB-WaVE latent space into five clusters.

**Figure S16:**
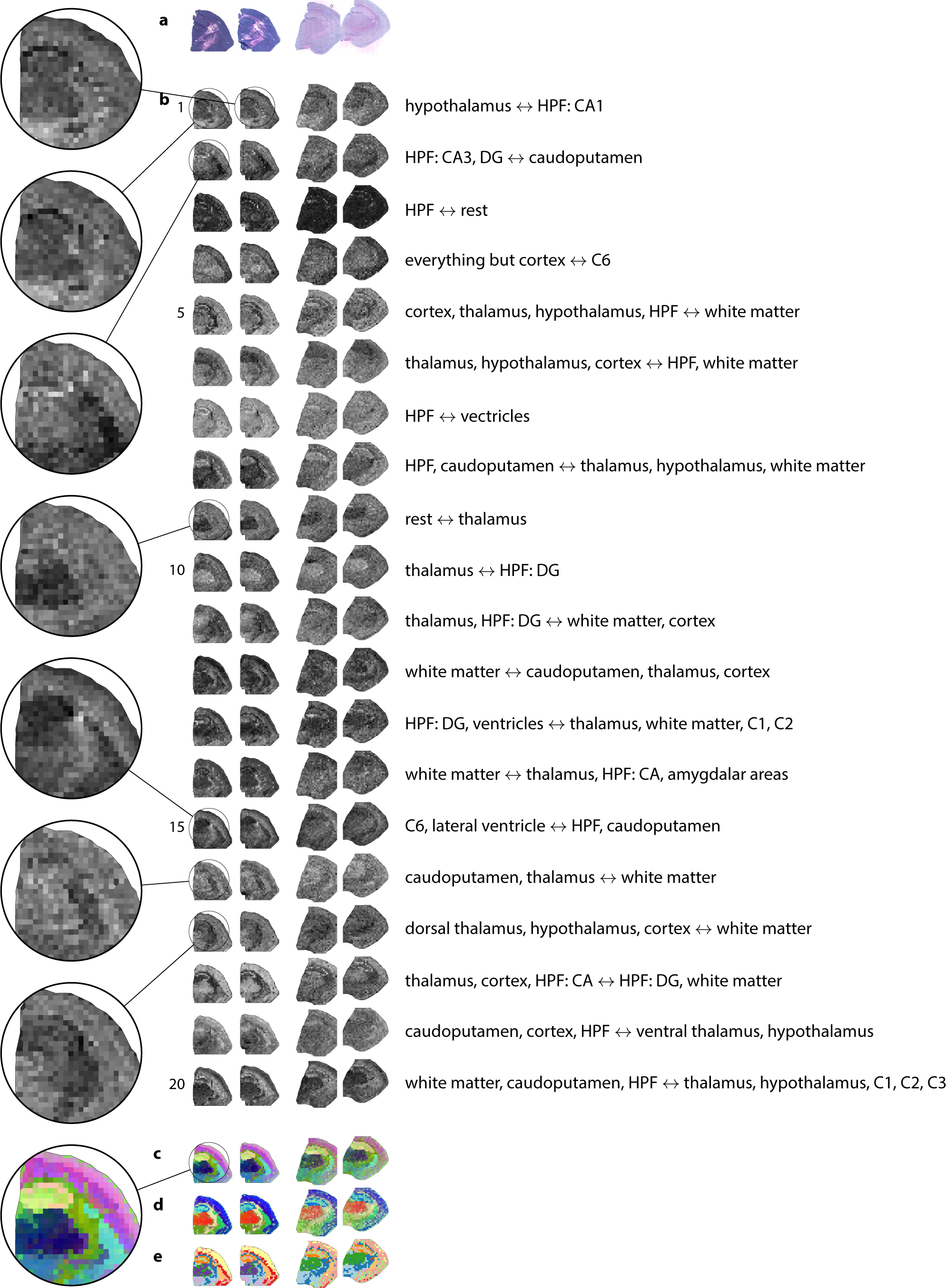
Results of analyzing mouse coronal brain sections using scVI. (**a**) H&E-stained microscopy images. (**b**) Latent space representation of scVI, and neuroanatomical axis corresponding to latent dimension. Note that the scVI latent dimensions are not sorted. (**c, d**) Summarization of scVI latent space, using t-SNE (**c**) or UMAP (**d**). (**e**) Hierarchical clustering of scVI latent space into twelve clusters. Abbreviations: see fig. 3.

**Figure S17:**
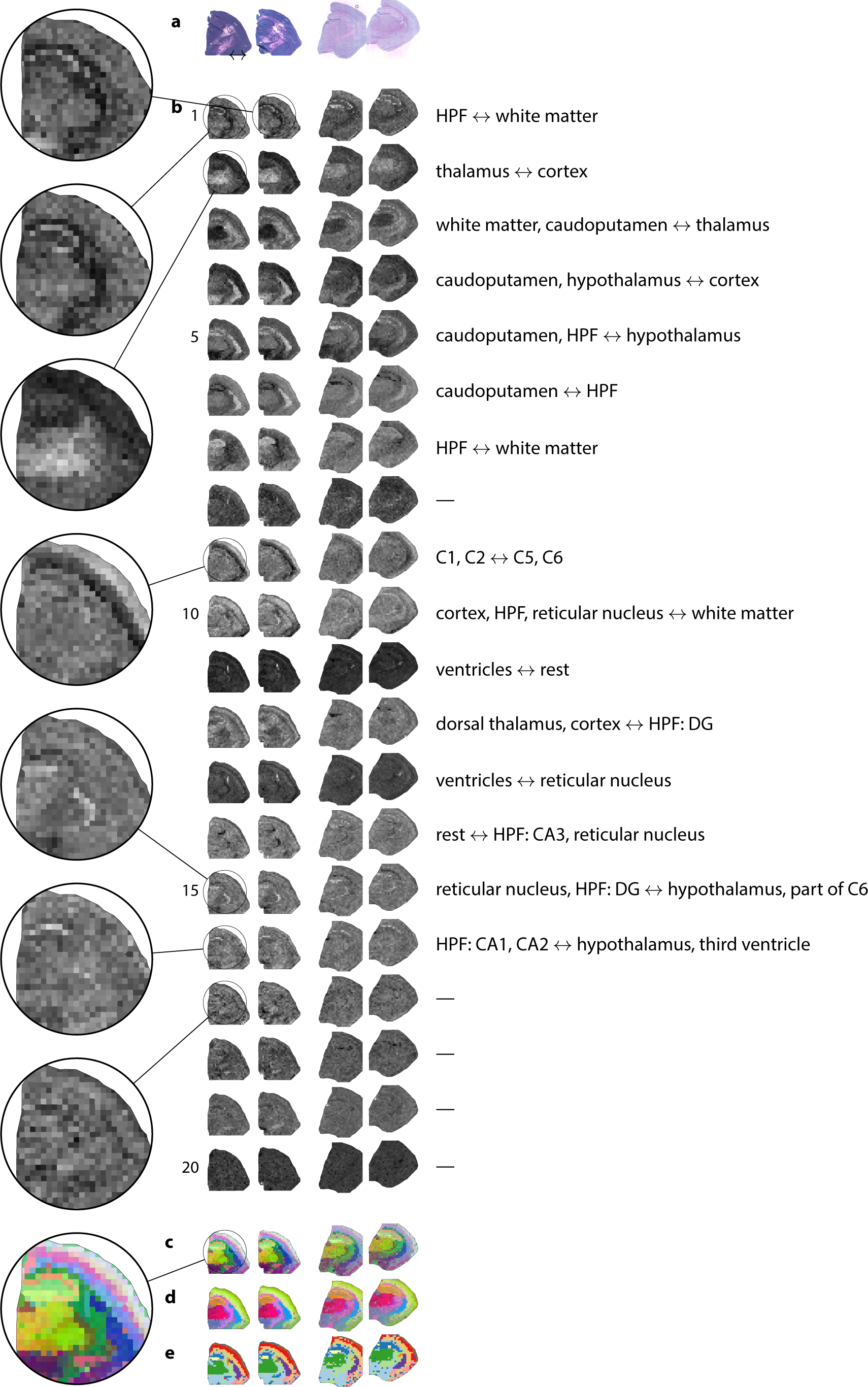
Results of analyzing mouse coronal brain sections using ZINB-WaVE. (**a**) H&E-stained microscopy images. (**b**) Latent space representation of ZINB-WaVE, and neuroanatomical axis corresponding to latent dimension. (**c, d**) Summarization of ZINB-WaVE latent space, using t-SNE (**c**) or UMAP (**d**). (**e**) Hierarchical clustering of ZINB-WaVE latent space into twelve clusters. Abbreviations: see fig. 3.

